# Allosteric priming of *E. coli* CheY by the flagellar motor protein FliM

**DOI:** 10.1101/781468

**Authors:** Paige Wheatley, Sayan Gupta, Alessandro Pandini, Yan Chen, Christopher J. Petzold, Corie Y. Ralston, David F. Blair, Shahid Khan

## Abstract

Phosphorylation of *Escherichia coli* CheY protein transduces chemoreceptor stimulation to a highly cooperative flagellar motor response. CheY binds to the N-terminal peptide of the FliM motor protein (FliM_N_). Constitutively active D13K-Y106W CheY has been an important tool for motor physiology. The crystal structures of CheY and CheY.FliM_N_ with and without D13K-Y106W have shown FliM_N_ bound CheY contains features of both active and inactive states. We used molecular dynamics (MD) simulations to characterize the CheY conformational landscape accessed by FliM_N_ and D13K-Y106W. Mutual information measures identified the central features of the long-range CheY allosteric network between D13K at the D57 phosphorylation site and Y/W106 at the FliM_N_ interface; namely the closure of the α4-β4 hinge and inward rotation of Y/W106 with W58. We used hydroxy-radical foot-printing with mass spectroscopy (XFMS) to track the solvent accessibility of these and other sidechains. The solution XFMS oxidation rate correlated with the solvent-accessible area of the crystal structures. The protection of allosteric relay sidechains reported by XFMS confirmed the intermediate conformation of the native CheY.FliM_N_ complex, the inactive state of free D13K-Y106W CheY and the MD-based network architecture. We extended the MD analysis to determine temporal coupling and energetics during activation. Coupled aromatic residue rotation was a graded rather than a binary switch with Y/W106 sidechain burial correlated with increased FliM_N_ affinity. Activation entrained CheY fold stabilization to FliM_N_ affinity. The CheY network could be partitioned into four dynamically coordinated community sectors. Residue substitutions mapped to sectors around D57 or the FliM_N_ interface according to phenotype. FliM_N_ increased sector size and interactions. These sectors fused between the substituted K13K-W106 residues to organize a tightly packed core and novel surfaces that may bind additional sites to explain the cooperative motor response. The community maps provide a more complete description of CheY priming than proposed thus far.

**Statement of Significance:** CheY affinity for FliM_N_, its binding target at the flagellar motor, is increased by phosphorylation to switch rotation sense. Atomistic simulations based on CheY and CheY.FliM_N_ crystal structures with and without the phospho-mimetic double substitution (D13K-Y106W) showed CheY compaction is entrained to increased FliM_N_ affinity. Burial of exposed aromatic sidechains drove compaction, as validated by tracking sidechain solvent accessibility with hydroxyl-radical foot-printing. The substitutions were localized at the phosphorylation pocket (D13K) and FliM_N_ interface (Y106W). Mutual information measures revealed these locations were allosterically coupled by a specialized conduit when the conformational landscape of FliM_N_-tethered CheY was modified by the substitutions. Novel surfaces stabilized by the conduit may bind additional motor sites, essential for the high cooperativity of the flagellar switch.

## Introduction

*Escherichia coli* CheY is a founding member of a bacterial response regulator superfamily that uses aspartate phosphorylation to regulate diverse signal relays (1, 2). The CheY β_5_α_5_ fold has structural homology with small eukaryotic signal-transducing proteins (3). CheY phosphorylation is essential for coupling chemoreceptor array states to motor response in bacterial chemotaxis. Previous studies of CheY have established it as a model for fundamental design principles in protein allostery (4).

Here, we study *E. coli* CheY binding to the FliM N-terminal peptide (FliM_N_) responsible for its initial interaction with the flagellar switch complex. Phospho-CheY produced within the chemoreceptor/CheA array diffuses to the flagellar motor within the flagellar basal body, interacting with its C-ring (a.k.a. the switch complex), a multi-subunit assembly consisting of the proteins FliG, FliM and FliN. The interaction increases clockwise {CW} rotation (5) in *E. coli*. In certain species such as *B. subtilis*, phospho-CheY exerts the opposite effect, stimulating CCW rather than CW rotation, but remains critical for chemotaxis. The affinity of non-phosphorylated *E. coli* CheY for FliM_N_ (ca. 600 μM) is 20x weaker than for phosphorylated CheY (6). The binding of activated CheY to flagellar switch complexes is non-cooperative (7), reflecting the interaction with FliM_N_. However, rotational bias displays a sigmoidal dependence on CheY concentration (Hill coefficient > 10.5) (8) implying highly cooperative action of the captured CheY molecules switching flagellar rotation with significant CheY occupancy of the FliM subunits, which number around 34 (9), for CW rotation to occur. Evidence that CheY interaction with the switch involves two binding sites, initial interaction with FliM_N,_ followed by a subsequent interaction of the FliM_N_ tethered CheY to FliN in *E. coli* (10) or the middle domain of FliM (FliM_M_) in *Thermatoga maritima* (11), provides a plausible mechanism for the high cooperativity. It has remained unclear whether the FliM_N_ tether facilitates the second-stage binding step only by increasing CheY local concentration, or whether structural changes also occur that prime CheY to bind FliN (**Figure 1**).

**Figure 1:**
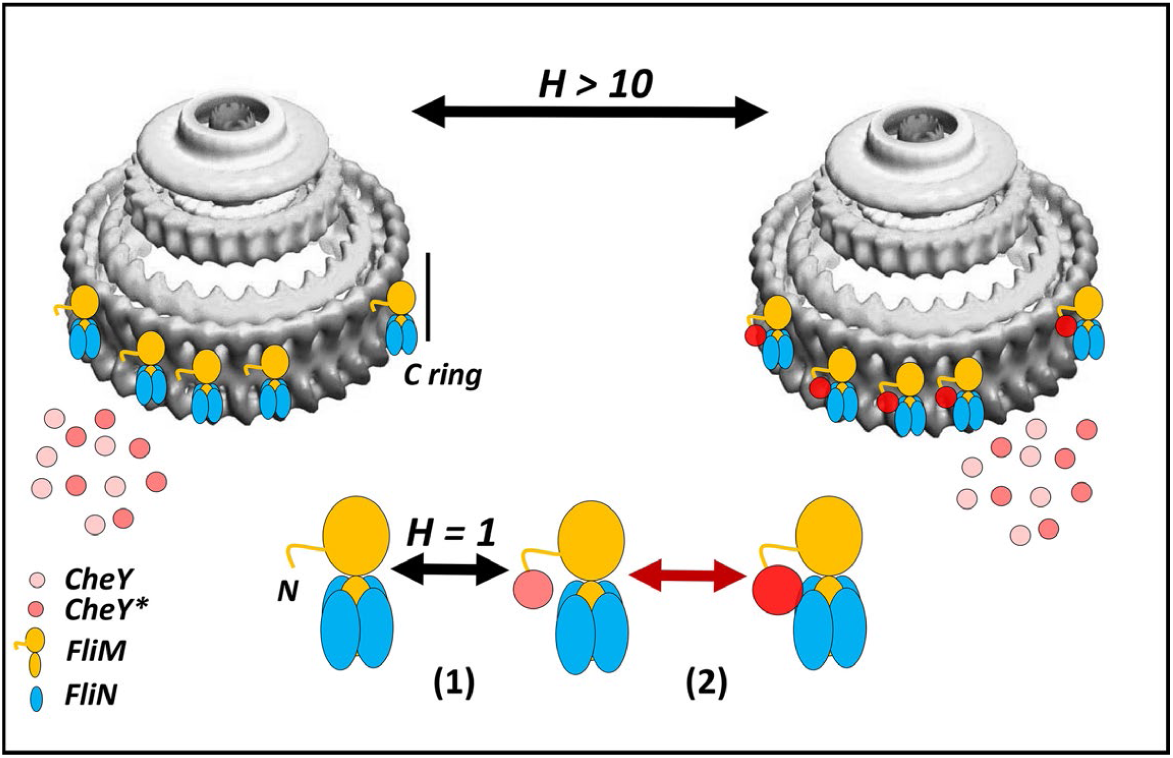
CheY interactions with the flagellar motor. CheY shade intensity and size denote activation state and FliN binding probability respectively. Binding of CheY* to isolated or in vivo flagellar basal bodies is not cooperative (H = 1), but the change in flagellar CW/CCW rotation bias is highly cooperative (H > 10) with CheY* concentration. 1^st^ stage (1) binding to FliM_N_ enables 2^nd^ stage (2) binding to FliN. The increased local concentration due to 1^st^ stage binding and the multiple FliN copies enhance 2^nd^ stage binding probability. Inactive CheY binds more weakly, reducing FliM_N_-tethered CheY binding events with FliN below the critical threshold for CW rotation. This study provides evidence for structural changes in CheY that may supplement increased local concentration for 2^nd^ stage binding.

Mutants that mimic aspects of the activation mechanism have been used to examine conformational changes in CheY and its interaction with FliM_N_ since the phosphoryl aspartate (D57∼PO_4_) is labile. Mutant *E. coli* CheY x-ray crystal structures, kinetic and behavioral assays (12-18) have outlined the structural changes during allosteric communication. W58 fluorescence is quenched when the adjacent D57 is phosphorylated. Suppressor mutations restore rotation bias in FliM chemotaxis (*che*) mutants. The activating substitutions D13K, Y106W are potent modulators of motor rotation bias in vivo (19-21) and focus of the present study.

Crystal structures of D13K-Y106W CheY alone and in complex with FliM_N_ showed bound FliM_N_ was required for the activated CheY conformation. They established CheY residues K91, Y106 and K119 as part of the FliM_N,_ binding surface and characterized their interactions (22). K91 and K119 formed salt bridges with FliM_N_. The W106 sidechain, exposed in CheY, moved in as FliM_N_ bound to switch K109 bonding interactions with T87, D57 and, via bound water, with D12 (22). The structure of the native CheY.FliM_N_ complex exhibited some features of inactive CheY and some features of the active D13K-Y106W CheY.FliM_N_ (23). Notably, the orientation of the Y106 sidechain matched that for W106 in the D13K-Y106W CheY.FliM_N_ complex. The “intermediate” conformation of the native CheY.FliM_N_ structure challenged two-state CheY allostery models that coupled Y/W106 rotamer state to T87 motions (24). CheY has high conformational plasticity as seen by the discrepancies between crystal structures of activated CheY proteins obtained by chemical modification versus residue substitutions (22, 25, 26). The coverage of the conformational landscape by crystal structures is too sparse to resolve the conformational trajectories for activation by phosphorylation or binding targets such as FliM_N_. Alteration of low-affinity binding interfaces, a common occurrence in signal-transducing phospho-relays by crystal packing contacts Is an additional concern for structures of protein complexes (27).

CheY conformational plasticity is not well-described by classical protein allostery concepts of “induced fit” (KNF) (28) or “conformational selection” (e.g. MWC (29)) but is accommodated by modern ideas of allostery (30) where protein-protein interactions between flexible partners have been described in terms of a folding funnel, where the funnel bottom has a “rugged” landscape with multiple minima (31). Accordingly, molecular dynamics (MD) simulations and solution measurements have supplemented the X-ray crystallography of free CheY structures. MD of free CheY examined the coupling between Y106 rotation and T87 movements triggered by hydrogen bond formation (32), showing that the β4-α4 loop is an important determinant of allosteric signaling affected by lysine acetylation (33) and extracted common design principles between CheY and other response regulators with correlation analyses (34, 35).

Here, we detail simulations and solution measurements to better understand the differences between the native and D13K-Y106W CheY crystal structures. We resolved the complex conformational landscapes by MD simulations with mutual information measures to determine the coupling between protein fragments. Protection experiments with XFMS (X-ray foot-printing with mass spectroscopy) (36, 37), a technique that probed sidechain solvent accessibility in contrast to deuterium exchange of backbone hydrogen atoms, supported the FliM_N_ requirement for D13K-Y106W CheY activation reported by the crystal structures, and the MD allosteric network model. XFMS has a more straight-forward physical rationale than fluorescence quenching for reporting sidechain motions over time-resolved windows and is not limited by the size of the protein assembly. Further analysis of the MD trajectories resolved multiple CheY Y106 rotamer states. Inward orientation was temporally coupled to stabilization of both the CheY fold and the FliM_N_ interface in the CheY.FliM_N_ complex, but not in CheY alone. The coupling increased in D13K-Y106WCheY.FliM_N_. The formation of a distinct module that orchestrates CheY dynamics to stabilize new surface topologies for possible second-stage binding to FliN was the signature of the fully activated D13K-Y106W CheY.FliM_N_ state

## Materials & Methods

### 1. Structure Preparation

Structures of *Escherichia coli* CheY (PDB ID: 3CHY. 1.7-angstrom resolution (38)) and complexes of native (PDB ID: 2B1J. 2.8 angstrom resolution (23)) and mutant (13DKY106W) CheY (PDB ID: 1U8T. 1.5 angstrom resolution (22)) with FliM_N_ were downloaded from Protein Data Bank. The 1U8T unit cell was a tetramer with 2 CheY and 2 CheY.FliM_N_ complexes. We generated the native CheY.FliM_N_ complex structure (1U8T_DY) by reverse mutagenesis (13 K->D, 106 Y->W) in silico to base the simulations on well-resolved atomic coordinates and eliminate the influence of crystal contacts. Missing atoms were added in Swiss-PDB viewer (www.expasy.org/spdbv); missing loop segments were completed with Modeller (https:/salilab.org/modeller). Mutant substitutions were made in Pymol (http://pymol.org) then energy minimized in Modeller.

Contact residues, surfaces and energies were extracted from the PDB files with the sub-routines (*ncont, pisa*) available in CCP4 version 7 (http://www.ccp4.ac.uk/). Comparison with experimental B-factors and geometrical analyses were performed with GROMACS version 4.5.7 (http://www.gromacs.org/). Solvent accessible surface area (SASA) was computed with the POPS server (https://mathbio.crick.ac.uk) (39). The orientation of the aromatic sidechains relative to the D57 C^α^ atom was measured in Pymol. It is specified by the angles θ and ω. The primary angle θ is given by the lines connecting the residue C^α^ (R-C^α^) to D57-C^α^ and its most distant sidechain heavy atom (OH (Y), CH2 (W, F). The angle ω specifies the orientation of the planar sidechain to the R-C^α^− D57-C^α^ line.

### 2. Molecular Simulations

#### (a) Molecular Dynamics

A set of 3 replicas of duration 1 μs each was generated for the mutant (1U8T) and native (1U8T_DY) complexes using GROMACS 2016.2 with Amber ff99sb*-ILDNP force-field (40). Another set of 3 replicas of 500 ns duration each was generated for the native CheY (3CHY). Each system was first solvated in an octahedral box with TIP3P water molecules with a minimal distance between protein and box boundaries of 12 Å. The box was then neutralized with Na^+^ ions. Solvation and ion addition were performed with the GROMACS preparation tools. A multistage equilibration protocol, modified from (41) as described in (42), was applied to all simulations to remove unfavourable contacts and provide a reliable starting point for the production runs including steepest descent and conjugate gradient energy minimisation with positional restraints (2000 kJ mol^-1^ nM^-2^) on protein atoms followed by a series of NVT MD simulations to progressively heat the system to 300 K and remove the positional restraints with a finally NPT simulation for 250 ps with restraints lowered to 250 kJ mol^-1^ nM^-2^. All the restraints were removed for the production runs at 300 K. In the NVT simulations temperature was controlled by the Berendsen thermostat with a coupling constant of 0.2 ps, while in the NPT simulations the V-rescale thermostat (43) was used with coupling constant of 0.1 ps and pressure was set to 1 bar with the Parrinello-Rahman barostat and coupling constant of 2 ps (44). Time steps of 2.0 fs with constraints on all the bonds were used. The particle mesh Ewald method was used to treat the long-range electrostatic interactions with the cut-off distances set at 12 Å. The MD runs reached stationary root mean square deviation (RMSF) values within 3 ns.

Collective motions were identified by PCA of the conformational ensembles. PCs were generated by diagonalization of the covariance matrix of C^α^ positions in GROMACS. The overlap (cumulative root mean square inner product) of the PCs between replicas (45)) and the PC dot product matrix was computed with the GROMACS g-anaeig function.

The conformational ensembles were clustered and mean structures representing the major clusters (n>5) computed with the GROMACS g-cluster function. The energy landscape was computed with PROPKA 3.0 (46). PROPKA calculates the free energy difference (ΔG) between the folded and unfolded states as the protein charge varies with pH (47). CheY has 37 ionizable groups (9D, 12E, 10K, 4R, 2Y) plus N and C termini that determine its net charge. The ΔG is computed from the perturbation of residue pK values by the protein environment; specifically the dielectric-dependant de-solvation penalty, backbone and sidechain hydrogen bonds and interactions with other charged residues. For the complexes, the ΔG was computed for the complex (ΔG_T_) as well as CheY alone with FliM_N_ removed (ΔG_CheY_). The ΔG_CheY_ was the free energy of the CheY fold. The interfacial energy ΔG_interface_ = ΔG_T_ - ΔG_CheY_

#### (b) tCONCOORD

tCONCOORD utilizes distance constraints based on the statistics of residue interactions in a crystal structure library (48, 49), to generate conformational ensembles from one crystal structure with solvent modelled as an implicit continuum. tCONCOORD runs compared conformational ensembles for native CheY (3CHY) with double-mutant CheY, extracted from the heterogenous 1U8T unit cell that contains structures both with and without FliM_N_. The random atom displacements were limited to 2 nm^3^, the iteration limit for the generation of a new structure was 500 and a threshold solvation score of 2.2 (48). Sets of 16^4^ = 65,536 equilibrium conformations with full atom detail were typically generated for each structure. The overlap between ensemble subsets was > 99% when the subset size was < 1/4 of this value (50).

The structure was rebuilt by random displacement of its atoms within limits, followed by an iterative correction to eliminate bond violations until a new structure that satisfies all bonding constraints is obtained. The process was repeated to generate an ensemble, with simulation parameters listed in Methods. Full atom detail is preserved, but the solvent is not modelled for increased computational speed. Instead, tCONCOORD has a solvation parameter that estimates the distance-dependent probability for a water molecule next to a particular atom to allow prediction of unstable hydrogen bonds (48) – solvent is “implicit”. The cost is reduced spatiotemporal detail due to loss of long-range and bound-water interactions.

### 3. Network Analysis

#### (a) Structural alphabet

Coordinated CheY motions were examined using mutual-information analysis. The mutual-information (nMI) matrix encodes correlations between conformational states of different parts of the protein backbone. The states are represented by a structural alphabet (SA). The SA (51), is a set of recurring four residue fragments encoding structural motifs derived from PDB structures. Fragments are assigned an SA designation according to backbone dihedral angles, allowing conformation to be specified in terms of a 1-dimensional string (51).

The mutual information *I*(*X*;*Y*) between two fragments (*X*) and (*Y*) is

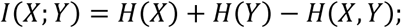

where *H*(*X,Y*) is the joint probability distribution

The normalized mutual information, *nMI*(*X*; *Y*) = (*I*(*X*; *Y*) − *ε*(*X*; *Y*))/(*H*(*X, Y*)); *H*(*X*) is a measure of the entropy Δ*S*(*X*) that is related to the number of microstates and their probability. *k*_*B*_ Is the Boltzmann constant. 

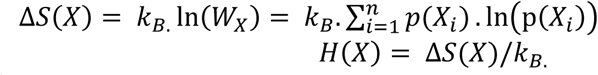

*ε*(*X*;*Y*) is the expected, finite-size error.

The finite-size error estimated as in earlier publications (e.g. (34, 52)) corrects for the effects of finite data and quantization on the probability distribution (53). The *nMI* couplings are detected as correlated changes in fragment dynamics, after spatial filtration to isolate long-range couplings (34). There is no need for discretisation and / or optimisation of parameters as the fragment set is pre-calculated. The fragments are represented as network nodes, with the connectivity (edges) between them representing their correlated dynamics over the MD trajectory.

#### (b) Eigenvector Analysis

Statistically significant correlations between columns were identified with GSATools (54) and recorded as a correlation matrix. The correlation matrix was used to generate a network model with the residues as nodes and the correlations as edges. The contribution of a node to the network was estimated by the eigenvector centrality, *E*, calculated directly from the correlation matrix: 

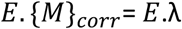

where {*M*}_*corr*_ is the correlation matrix, the *λ* is the eigenvalue

The nMI contribution of local fragment motions was computed for the top PCs and superimposed on their RMSF profiles to evaluate the mechanical behavior of the network nodes in driving collective motions.

Ensemble conformations and MD runs were averaged for computation of the nMI between fragment positions, with > 2σ thresholds for selected top couplings. Pearson’s correlations were used for comparison. Significance limits were set in GSATools.

#### (c) Community Analysis

The Girvan–Newman algorithm (55) was used to identify community structure. Then the network was collapsed into a simplified graph with one node per community, where the node size is proportional to the number of residues. Edge weights represent the number of nMI couplings between communities (56). Community analysis of correlation networks identifies relatively independent communities that behave as semi-rigid bodies. Graphs were constructed with the *igraph* library (57) in R (https://cran.r-project.org/web/packages/igraph/) and visualized in Cytoscape (http://www.cytoscape.org/).

### 4. Overexpression and purification of CheY proteins

*E. coli* strain BL21/DE3 with CheY-pET21b plasmid (10) was grown in 15 mL LB + 100 ug/mL ampicillin at 32°C for 15-18 hours. The culture was diluted 1:100 into 1 L LB + 100 ug/mL ampicillin + 250 uM IPTG and grown at 32°C until OD_600_ 0.5-0.7. Cells were collected by centrifugation at 5000xg for 20 minutes at 4°C. The cell pellet was resuspended in 8 mL lysis buffer (1 mg/ml lysozyme, 1 mM PMSF, 50 mM NaH_2_PO_4_, 300 mM NaCl, 10 mM imidazole, pH 8) and rocked on ice for 2 hours. 5 ug/mL DNAse I and 2.5 mM MgCl_2_ were added and the lysed solution was rocked on ice for an additional 20 minutes, or until the lysate was no longer viscous. Lysed samples were sonicated for 30 pulses (Branson Sonifier 250, duty cycle 50, output 3) to completely lyse cells. The lysate was centrifuged at 10,000g for 30 minutes at 4°C to remove cellular debris from the His-tagged CheY protein. The lysate supernatant was combined with 4 mL Ni-sepharose beads and mixed gently at 4°C for 1 hour. The CheY-HIS-Ni-sepharose beads were washed twice with 10 mL 50 mM NaH_2_PO4, 300 mM NaCl, 20 mM imidazole, pH 8; twice with 10 mL 50 mM NaH2PO4, 300 mM NaCl, 30 mM imidazole, pH 8; and twice with 10 mL 50 mM NaH_2_PO4, 300 mM NaCl, 40 mM imidazole, pH 8. CheY protein was washed off the beads 7-8 times with 1mL 50 mM NaH_2_PO_4_, 300 mM NaCl, 250 mM imidazole, pH 8. 4 mL of purified CheY protein was dialyzed into XF (X-ray foot-printing) buffer (10 mM potassium phosphate buffer (pH 7.2), 100 mM NaCl, and 10 mM MgCl2) at 4°C.

Purified CheY, FliM_N_.CheY, CheY*, and FliM_N_.CheY* proteins were analyzed by fast protein liquid chromatography (FPLC) on an AKTA Superdex-75 10/300 GL column in X-ray foot-printing buffer at 22°C (conductivity 28.9 mS/cm). Carbonic anhydrase (29 kDa) and ribonuclease A (13.6 kDa) were also analyzed on the same Superdex-75 column in X-ray foot-printing buffer as molecular weight standards **(Supporting Information Figure S1)**.

The foot-printing experiments used FliM_N._CheY and FliM_N._CheY* fusion proteins in addition to CheY and CheY*. The experimental fusion protein data was compared against simulations based on crystal structures of the corresponding CheY.FliM_N_ complexes. The FliM_N._CheY fusion interacts with FliN (10) and is more potent than CheY alone in potentiation of CW rotation (P. Wheatley, unpublished). 3D models of the FliM_N_.CheY fusions were obtained with the I-Tasser suite (58). I-Tasser uses threading and template-matching to determine secondary structure segments. Tertiary 3D topology is generated by *ab initio* folding of undetermined regions by replica-exchange Monte-Carlo dynamics and refined to remove steric clashes and optimize hydrogen bonding. The final models are assigned a confidence score (cs = -5.0 -> 1.0) based on multi-parametric comparison against Protein Data Bank structures. In all top 5 models, FliM_N_ was docked in the location seen in the crystal structures of the CheY.FliM_N_ complexes. The top model had cs = -1.08, RMSF = 7.2+4.2 angstroms (against CheY, FliM_N_ crystal structures).

### 5. X-Ray Foot-printing (XF)

Protein samples (CheY, FliM_N_.CheY, CheY*and FliM_N_CheY* were prepared in 10 mM potassium phosphate buffer (pH 7.2), 100 mM NaCl, and 10 mM MgCl_2_. Exposure range was determined empirically by adding Alexa488 to protein solutions as previously described (59). Sample irradiation was conducted without Alexa488 dye using a microfluidic set-up with 100 mm and 200 mm ID tubing in combination with a syringe pump as previously described (60). After exposure at ALS beamline 3.2.1, samples were immediately quenched with methionine amide to stop the secondary oxidations and stored at -80 °C for LCMS analysis.

The oxidized fraction, F, for a single residue modification was given by the equation 

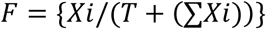

where *XX* is the oxidized residue abundance of one of the monitored residues in a trypsinized peptide and *T* is the unoxidized peptide.

Best fit first-order rates were calculated in Sigmaplot version 12. Protection factors (PFs) were calculated as the ratio of the intrinsic residue reactivity over its foot-printing rate (61). Its logarithm (log(PF)) was proportional to the SASA. The relation assumes that the foot-printing rate was related to the activation energy associated with the accessibility of the side-chain to hydroxy radicals and the initial step of hydrogen abstraction It empirically gave the best-fit for proteolyzed peptides on a model data set, extended here to single residues (61).

### 6. Mass Spectrometry (MS) Analysis

X-Ray exposed protein samples were digested by Trypsin and the resulted peptide samples were analyzed in an Agilent 6550 iFunnel Q-TOF mass spectrometer coupled to an Agilent 1290 LC system (Agilent Technologies, Santa Clara, CA). Approximately 10 pmol of peptides were loaded onto the Ascentis Peptides ES-C18 column (2.1 mm x 100 mm, 2.7 μm particle size; Sigma-Aldrich, St. Louis, MO) at 0.400 mL/min flow rate and were eluted with the following gradient: initial conditions were 95% solvent A (0.1% formic acid), 5% solvent B (99.9% acetonitrile, 0.1% formic acid). Solvent B was increased to 35% over 5.5 min, and was then increased to 80% over 1 min, and held for 3.5 min at a flow rate of 0.6 mL/min, followed by a ramp back down to 5% B over 0.5 min where it was held for 2 min to re-equilibrate the column to original conditions. Peptides were introduced to the mass spectrometer from the LC using a Jet Stream source (Agilent Technologies) operating in positive-ion mode (3,500 V). Source parameters employed gas temp (250°C), drying gas (14 L/min), nebulizer (35 psig), sheath gas temp (250°C), sheath gas flow (11 L/min), VCap (3,500 V), fragmentor (180 V), OCT 1 RF Vpp (750 V). The data were acquired with Agilent MassHunter Workstation Software B.06.01 operating in either full MS mode or Auto MS/MS mode whereby the 20 most intense ions (charge states, 2–5) within 300-1,400 m/z mass range above a threshold of 1,500 counts were selected for MS/MS analysis. CheY native and oxidized peptides were identified by searching MS/MS data against *E. coli* protein database with Mascot search engine version 2.3.02 (Matrix Science). Based on the information of accurate peptide m/z value and retention time, the peptide precursor peak intensities were measured in MassHunter quantitative analysis software.

## Results

We analyzed conformational ensembles generated by MD to identify dynamic changes in CheY architecture, using loops and residues implicated in the allosteric relay (see Introduction) as markers. We used XFMS protection experiments to relate the crystal structures to the conformation landscape in solution and test the dynamics predicted by the MD simulations (**Figure 2A**).

**Figure 2:**
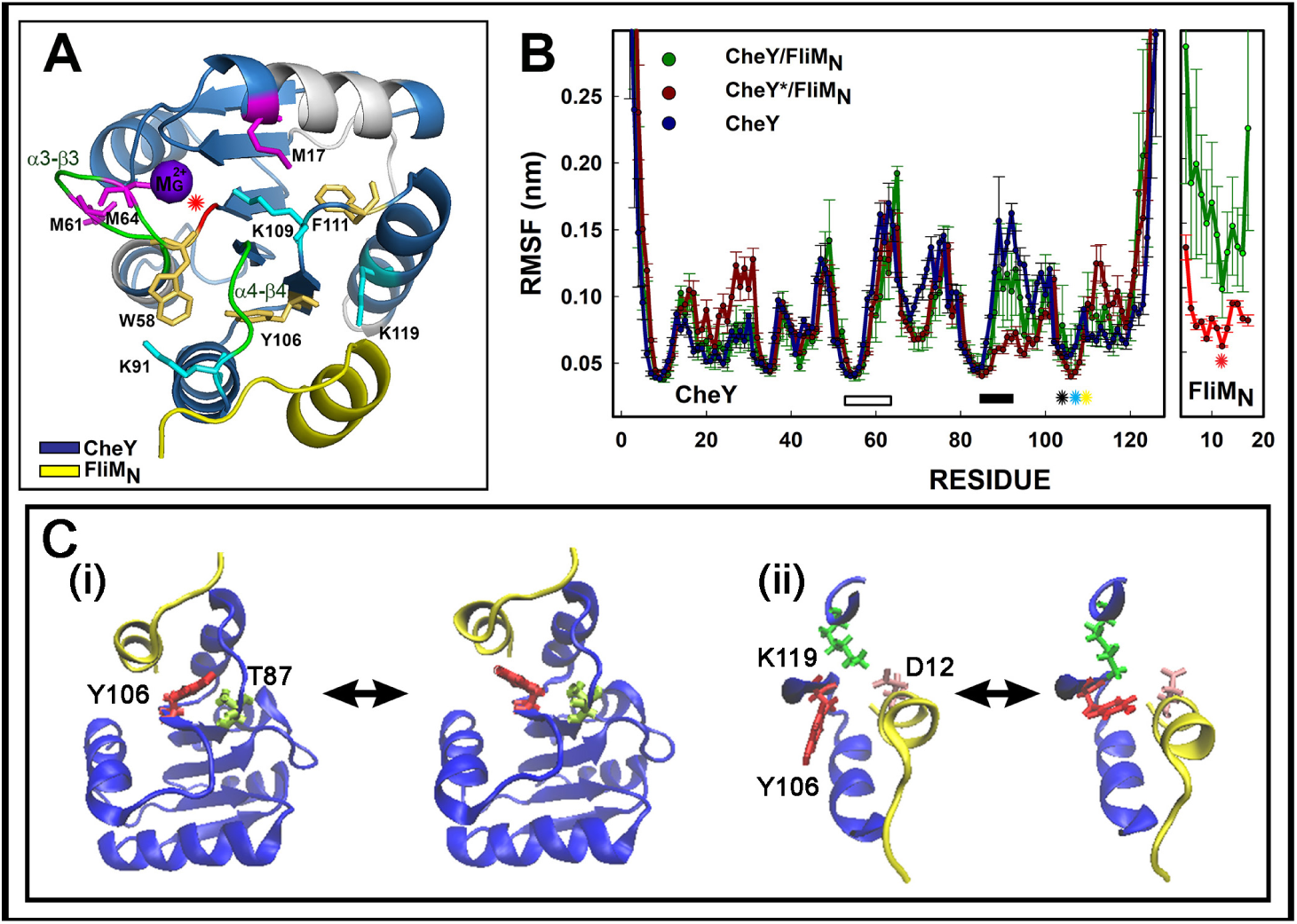
Dynamics of CheY-FliM_N_ association. **A.** Structure of CheY in complex with FliM_N_ (2B1J-AC.pdb). Colors indicate FliM_N_ (yellow), tryptic CheY fragments (blue), allosteric relay loops (green), sidechains (M(magenta), K (cyan), Y,W,F (gold). D57 C^α^ (red asterisk), Mg^2+^ (magenta), **B.** MD RMSF profiles for the combined replica trajectories for the three structures analyzed in this study. Bars mark CheY loops α3-β3 (white) and α4-β4 (black). Asterisks mark residues Y/W106 (black), K109 (cyan) and F111 (yellow). FliM_N_ residue D12 (red asterisk) forms a salt-bridge with CheY K119. **C.** Snapshots of CheY (blue) Y106 (red) transitions in 1U8T_DY coupled to internal and interfacial residues. FliM_N_ (yellow). (i) T87 (lime). Supporting Information Video S2. (ii) K119 (green), FliM_N_ D12 (pink). Supporting Information Video S4.

### 1. CheY activating residue substitutions D13K/Y106W stabilize FliM_N_ association

Three MD replica runs each was performed for the native CheY structure (3CHY.pdb(38)), the activated D13K-Y106WCheY in complex with N-terminal FliM peptide (FliM_N_) and alone (1U8T.pdb(22)), and a complex of native (non-activated) CheY with FliM_N_ engineered in silico from 1U8T.pdb (Methods). The crystal structures showed residue Y106 was in the OUT conformation in CheY (3CHY) and D13K-Y106WCheY (1U8T), but in the IN conformation in CheY.FliM_N_ (2B1J) and D13K-Y106WCheY.FliM_N_ (1U8T). The Y/W106 rotamer state was correlated with the orientation of the W58 and F111 sidechains (**Supporting Information Figure S2A**). The engineered complex was used instead of the crystal structure (2B1J.pdb) (23) since the latter, in addition to the lower resolution, had a systematic bias in its RMSF profile from the N- to C-terminus that may be due to mosaicity in the crystal consistent with increased CheY-FliM_N_ interfacial dynamics (**Supporting Information Figure S2B**).

Henceforth, the double substitution D13K-Y106W CheY will be referred to as “CheY*”; the 1U8T_DY CheY. FliM_N_ as the “native CheY complex” and CheY* FliM_N_ as the “mutant CheY complex”. The root-mean-square fluctuation (C^α^ RMSF) profile for each structure, averaged over three 1 μs runs, are shown in **Figure 2B.** The MD excluded the first three residues (M_1_GD_3_) of the FliM_N_sequence (M_1_GDSILSQAEIDALL_16_) as these were not resolved in the 1U8T structure. The profiles are compared with B-factors for the X-ray structures in Figure S2. The B-factors were high relative to the MD-derived RMSF’s, particularly in loop regions, reflecting conformational heterogeneity of these segments in the crystals.

The 3CHY MD trajectories revealed transitions of Y106 between the OUT and IN states, consistent with electron density observed for both states in the crystal structure. FliM_N_ secondary structure, the CheYK119-FliM_N_D12 salt-bridge and Y106/W106 rotamer state were conserved between the 2B1J and 1U8T crystal structures. However, the raw MD trajectories of the complexes showed FliM_N_ had higher mean C^α^ RMSF values when CheY was wild type than when it carried the activating substitutions **(Supporting Information Videos S1, S2, S3).** This difference was due to association/dissociation of the FliM_N_ N and C termini from native CheY. In CheY*.FliM_N_ trajectories the peptide centre was tethered by the CheYK119-FliM_N_D12 salt bridge. W106 was locked IN and part of the segment with the lowest C^α^ RMSF together with K109 and F111. In CheY.FliM_N_ trajectories, OUT excursions of Y106 cleaved this salt-bridge and weakened interfacial attachments **(Figure 2C, Supporting Information Video S4)**. Thus, the MD confirmed the suggestion from the CheY.FliM_N_ crystal structure that its FliM_N_ interface was labile.

### 2. Two loops control CheY network dynamics

We measured the mutual information between local protein fragments to formalize the structural basis for the conformational dynamics. The 3D dynamics were extracted as 1D strings with the SA. Mutual information provided a quantitative readout, represented as a network, of the dynamics that drove the conformational transitions. First, we identified the central nodes in the CheY global network with the highest connectivity (**Figure 3A, B**). In vector notation, the overall connectivity of a given fragment is reported by its eigenvector centrality (“centrality”). The central CheY nodes were the loops β3-α3 (D_57_WNMPNMDG) and β4-α4 (T_87_AEAKK). A third prominent node just below the 1σ threshold was the short β_5_-strand (Y_106_VVKP).

**Figure 3:**
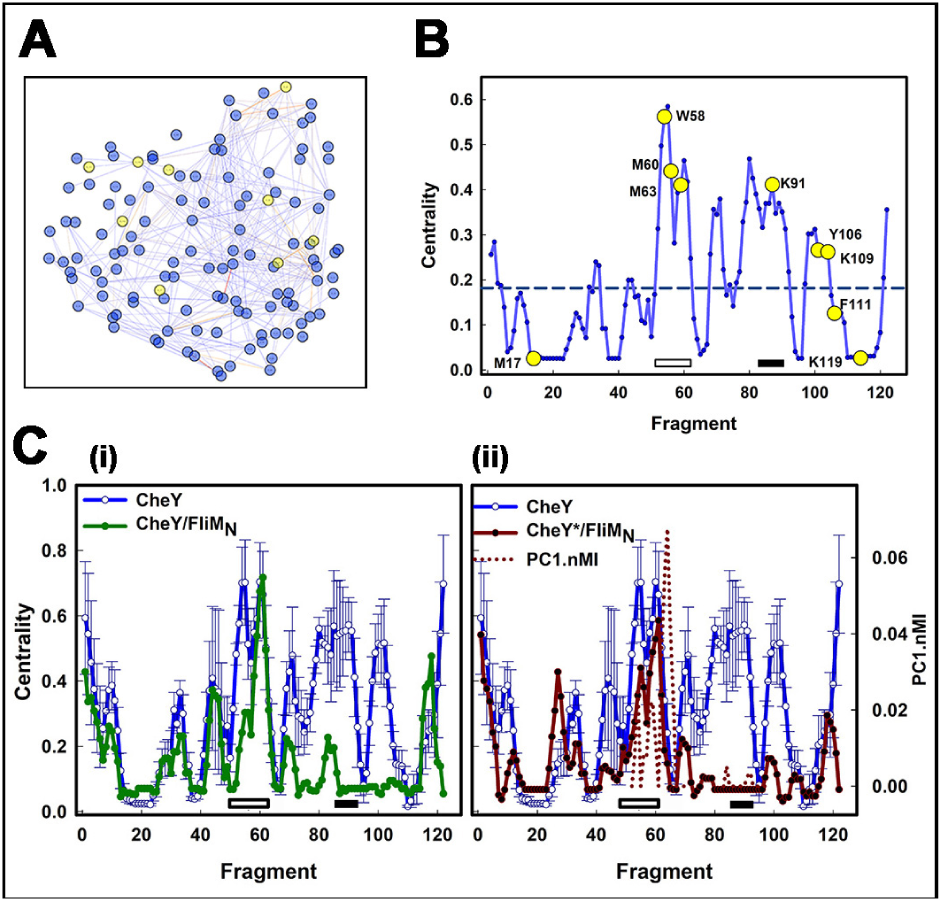
CheY network dynamics. **A.** The global network has nodes (residue fragments) and edges (mutual information weighted node interactions). **B.** The connectivity of the network is determined by the eigenvector centrality, a measure of the influence of individual nodes in the network. Nodes containing residues that are part of the allosteric relay (W58, K91, Y106, K109, F111) have high scores in the CheY network. In A, B, these residues and control residues monitored by XFMS (M17, M60, M63) are highlighted (yellow circles). **C.** Centrality profiles of the complexes. (**(i)** CheY/FliM_N_ (green). **(ii)** CheY*.FliM_N_ (red)) compared with the native CheY profile (mean ± s.e; blue lines). The dotted line ((ii) red) plots the mutual information between the local loop fragment dynamics and collective PC1 motions. Complex formation reduced the centrality of the β4-α4 loop that together with the β3-α3 loop formed central nodes in the CheY network. Activating mutations eliminated the β4-α4 loop as a node but did not alter the contribution of the β3-α3 loop in CheY*.FliM_N_. Horizontal bars indicate α3-β3 (white) and α4-β4 (black) loops as in Figure 2B.

Interpretation of differences between CheY and CheY* crystal structures based on isolated landmarks (for example, Y106 rotamer state (IN/OUT in CheY (3CHY.pdb) (38)) versus W106 (OUT in CheY*(22)) is complicated by the CheY conformational plasticity. Therefore, we used tCONCOORD, a computationally inexpensive method to generate conformational ensembles for comparison of the CheY and CheY* conformational landscapes. Analysis of the tCONCOORD ensembles showed the central network nodes remained unchanged, with both CheY Y106 and CheY* W106 sidechains restricted to a limited OUT-orientation range (**Supporting Information Figure S3**).

We next examined the CheY and CheY* complexes with FliM_N_ (**Figure 3C, D**). We split the CheY ensemble into four sub-populations to assess the significance of differences observed between it and the complexes. The network connectivity, as formalized by centrality plots, showed significant changes in the complexes relative to the CheY protein alone. There was a dramatic reduction in the centrality of loop β4-α4 and associated β-strand *_106_VVKP (* = Y (CheY.FliM_N_), W (CheY*.FliM_N_) at the FliM binding surface. Their roles as network nodes were reduced in CheY.FliM_N_ and abolished completely in CheY*.FliM_N_. This trend contrasted with the conservation of these nodes for CheY*.

### 3. Immobilization of the α4-β4 loop modulates CheY collective motions

We used Principal Component Analysis to characterize CheY collective motions and their modulation by FliM_N_ binding and the activating substitutions. The Principal Components (PCs) are derived from the atomic-coordinate covariance matrix and describe C^α^ backbone movements, ranked according to the amplitude of the structural variation they explain. The collective motions were described well by the first few PCs, as found for other proteins. The first three principal components (PCI-PC2-PC3) accounted for > 60% of all motions in each case (**Supporting Information Figure S4A, B**). These three PCs comprise bending and twisting modes organized around the β-sheet core. A core sub-population of CheY conformations was observed in MD trajectories generated by all three structures. When CheY is in complex with FliM_N_, new sub-populations comparable in size to the core were generated. These were distinct in the CheY.FliM_N_ and CheY*.FliM_N_ complexes. Thus, new conformational ensembles are accessed upon binding of FliM_N_, with the potential to produce binding surfaces for additional targets.

Loops act as hinge elements for collective motions. Their mechanics give insight into the modules they control (31). We computed loop β3-α3 and β4-α4 hinge flexibility by mapping their RMSF onto the PC1 that accounts for > 40 % of the total amplitude of the PC motions (Supporting Information Figure S4B). Flexibility scaled with the magnitude of the loop RMSFs relative to the mean PC1 RMSF. We computed hinge contribution to the PC1 as the nMI between PC1 variance and the local loop fragment dynamics (**Supporting Information Figure S4C**). The long β3-α3 loop partitioned into two segments. The short D_57_WN and the adjacent M_60_PNMDG loop segments behaved as rigid (low RMSF) and flexible (high RMSF) hinges respectively to control native CheY PC1 dynamics. In the CheY*-FliM_N_ complex, the β3-α3 loop hinge was retained, but with inverted flexibility of the two segments. The transition for loop β4-α4 was more dramatic from a flexible hinge in native CheY to a closed hinge that acted as a rigid lever arm in CheY*-FliM_N_. The reduced flexibility decreased β4-α4 loop centrality and influence on PC1 motions,

### 4. Protection experiments support the “intermediate” CheY.FliM_N_ structure and the MD allosteric network

We studied homogenous solutions of CheY and FliM_N_-CheY fusion proteins to measure the changes predicted by the crystal structures and the MD network model. The fusions were critical since the affinity of FliM for CheY is weak and that for the inactive protein even weaker (Introduction). The structures showed that **(i)** Aromatic sidechain internalization in CheY was entrained to FliM_N_ attachment, and **(ii)** The configurations of free CheY with or without the D13K-Y106W substitutions were similar. The MD revealed **(iii)** FliM_N_ attachment was more labile in the native versus D13K-Y106W complexes, and **(iv)** generated a network model to discriminate between CheY fragments that changed upon activation from those that did not. These predictions were assessed by comparing the sidechain solvent accessibility of allosteric relay residues Y106, W58, K91, K109, F111 and K119 by hydroxyl radical foot-printing in the native and D13K-Y106W CheY proteins, and their FliM_N_-fusion constructs. The control residues predicted not to change during activation were the β3-α3 loop residues M60 and M63 in proximity to W58, and the M17 in proximity to D/K13.

Aromatic residues have high intrinsic sidechain reactivities with hydroxyl radicals, exceeded only by methionine and cysteine (absent from *E. coli* CheY) followed by the alkaline sidechains. Tryptic digestion partitioned CheY into six separated peptides that were distinguished by mass spectroscopy (MS) based on their characteristic m/z ratio, allowing oxidation of these residues to be monitored. Dose-response curves were generated for each of the four constructs (CheY, CheY*, CheY-FliM_N_, CheY*-FliM_N_). For each residue examined, the curves from two independent experiments were pooled (**Supporting Information Figure S5**).

CheY residues of the allosteric relay at the FliM_N_ interface and distant from it were designated “interface” and “core” residues respectively. The oxidation of the interfacial residues (K119, Y/W106, K91) was reduced in the complexes (**Figure 4 A. Ii-iii**). Importantly, oxidation of the core residues also decreased with complex formation (**Figure 4 A. Ci-iii)**. In contrast, there was no significant difference between oxidation rates for β3-α3 loop control residues M60, M63 in the fusion proteins versus the free proteins, while the oxidation of the control M17 in the fusions was comparable or greater than in the corresponding free CheY proteins (**Figure 4B**).

**Figure 4:**
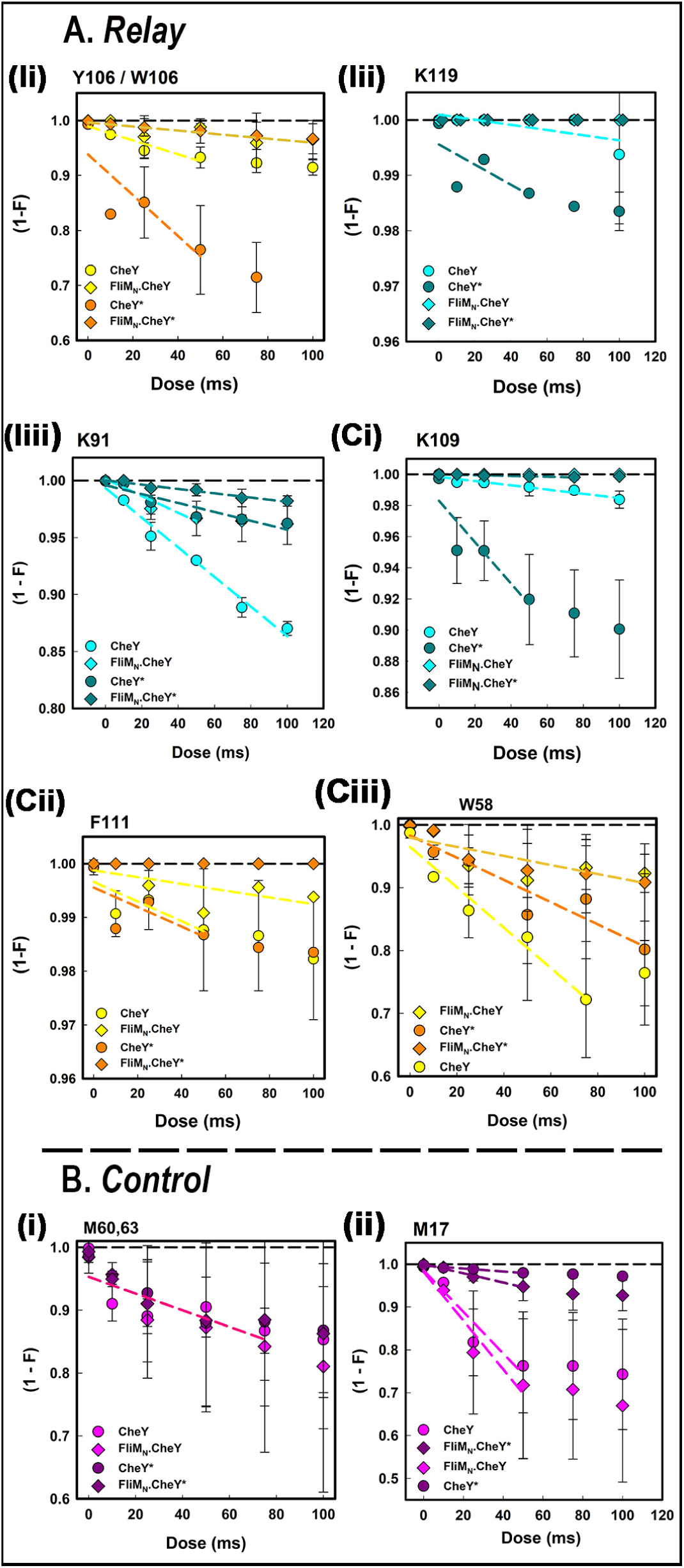
XFMS Measurements. Dose-response curves for **A**. Relay. Interfacial residues **(Ii)** Y/106W. **(Iii)** K119**. (Iiii)** K91. Core residues **(Ci)** K109. **(Cii)** F111 **(Ciii)** W58. **B**. Control residues. **(i)** M60, M63. **(ii)** M17. Initial rates (dashed lines) were obtained from least-squares linear regression of the decrease in the un-oxidized fraction with dose.

Protection factors (PFs) were computed from the initial rates from the single residue dose-response curves following protocols established by the study of 24 peptides from 3 globular model proteins (61), with intrinsic reactivities mostly determined thus far from measurements on small peptides (62). We first evaluated the agreement between solvent accessibility reported by the XFMS measurements and the crystal structures. Protection factors read-out the solvent-accessible surface area (SASA), with some caveats (61), The log(PF)s were plotted against the residue SASA in the crystal structures. The overall correlation was comparable to published values for the peptide correlations for the model proteins (61), indicating that the changes in the dose-response plots for the monitored residues are due, in large part, to non-polar bulk solvent accessibility changes (**Figure 5A**). Outliers (M17, K109, F111) were restricted to a small CheY protein volume in the structures (**Supporting Information Figure S6**). The crystal structures may not reflect the solution conformation of this local region, but bonding interactions may also contribute (Supporting Information Figure S6 legend). The correlation improved markedly (0.60 -> 0.86), without further correction, if the outliers were excluded.

**Figure 5.**
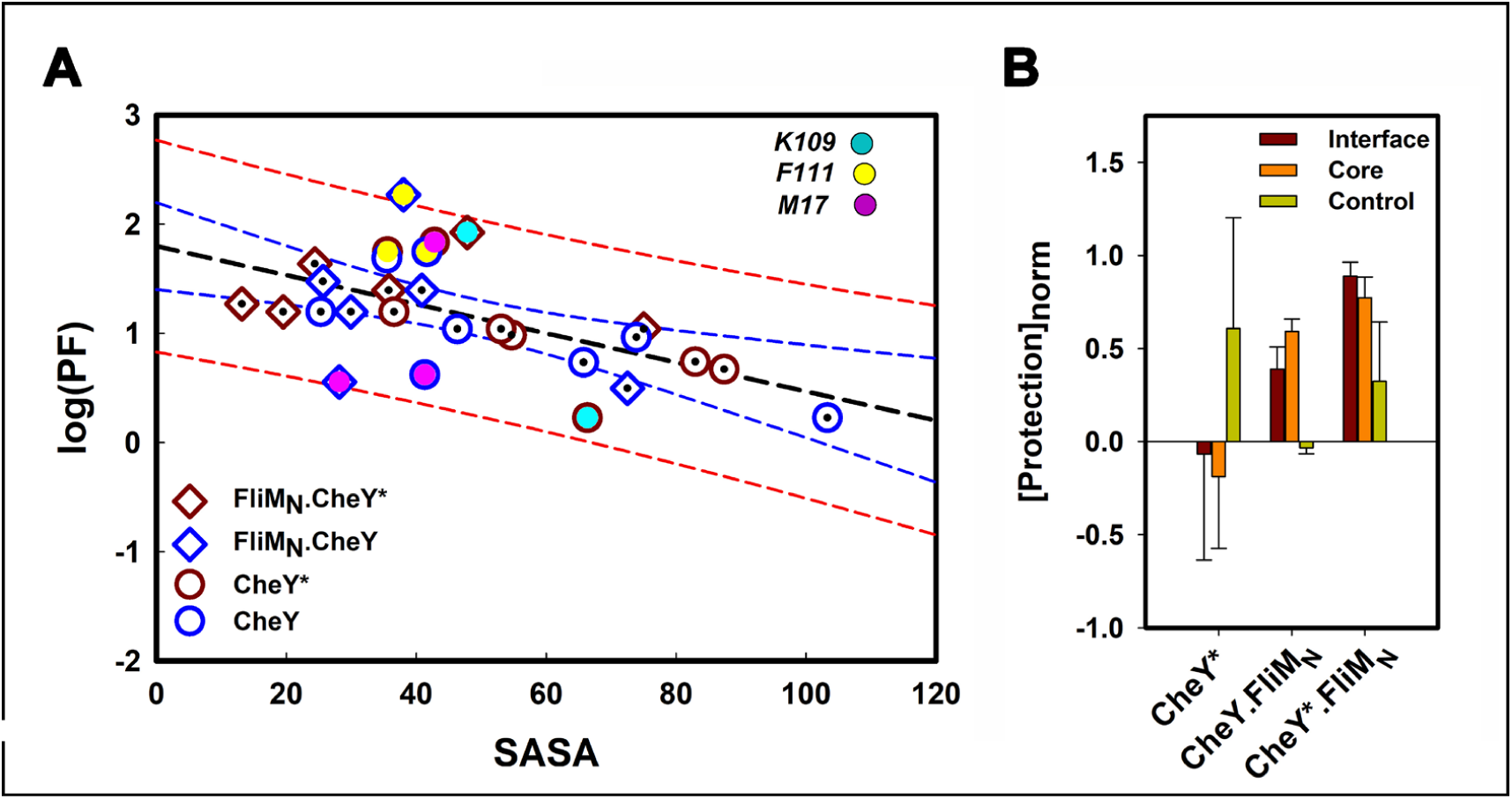
Single residue oxidations related to SASA. **A.** Log (PF)s plotted against the side-chain solvent accessible surface area (SASA) calculated from the crystal structures. Pearson correlation coefficients: 0.86 (minus M17 (rose), K109 (cyan). See text). Overall = 0.60 {CheY= (-)0.76; CheY*= (-)0.70; FliM_N_.CheY= (-)0.54;FliM_N_.CheY*=(-)0.12}. Best-fit (black dashed line), 95% confidence limit (blue lines), 95% prediction limit (red lines). **B.** Protection of interfacial (K119, Y/W106, K91), core (F111, K109, W58) and control (M17, M60, M63) residues in CheY*, CheY.FliM_N_, CheY*FliM_N_ relative to their protection in CheY. {Protection}_norm_ = Log {PF/PF_CheY_}. Positive values indicate increased protection.

The PFs for CheY*, CheY.FliM_N_ and CheY*.FliM_N_ were then normalized for each residue against the value obtained for CheY (**Figure 5B**). The normalized (log(PF)s) provided a quantitative measure for the increase for both interfacial and core residues in the CheY.FliM_N_ and CheY*.FliM_N_ fusions relative to the values for CheY. These residues were significantly more protected in CheY*.FliM_N_ than CheY.FliM_N._ In contrast, the protection of the control residues in the fusions (CheY.FliM_N_, CheY*.FliM_N_) did not differ significantly from that measured for CheY. The normalized PFs showed no significant difference in protection for interfacial, core or control residues in CheY* relative to CheY.

The protection profiles showed that solvent accessibility for the allosteric relay residues decreased in the order CheY < CheY.FliM_N_ < CheY*.FliM_N_. The control residues either showed no changes or the opposite trend. Changes in the solvent accessibility of CheY* relative to CheY were not significant. Thus, in conclusion, the XFMS experiments validated the main predictions of the crystal structures and the conformational ensembles generated from them.

### 5. Energetics of CheY stabilization by FliM_N_ and D13K/Y106W residue substitutions

The XFMS measurements correlated population shifts in selected residue positions to each other and with shifts in the crystal structures. The temporal couplings between these shifts could only be studied with MD. We next analyzed the MD trajectories to extract this information.

First, we examined the temporal coupling between the electrostatic stabilization of the interface and the CheY fold with the rotational states of residue Y106 (106W in CheY*.FliM_N_). CheY*.FliM_N_ 106W sidechain was locked IN (Supporting Information Video S3). In contrast, Y106 in CheY (Supporting Information Video S1) and CheY.FliM_N_ (Supporting Information Video S2) made frequent OUT <-> IN excursions. Dwell times in the Y106 rotamer states measured from the raw CheY trajectories were 107±34 ns (OUT) and 15±4 ns (IN)). The CheY.FliM_N_ Y106 sidechain was predominantly in the IN orientation, with mean dwell time 239±123 ns, 15-fold greater than for free CheY. The conformational ensembles in the MD trajectories were clustered based on the C^α^ backbone dynamics {RMSF} The major clusters represented distinct backbone conformational states accessed during the MD runs. The average structures for these clusters were compared to each other and the crystal structures with PROPKA. The mean ΔG values at pH 7.0 were CheY (−4.8±1.0 (n=7)) < CheY.FliM_N_ (−5.8±1.6 (n=4)) < CheY*.FliM_N_ (−9.9±2.2 (n=3)). All CheY clusters had Y106 in the OUT orientation (θ = 126.7±3.8_°_) indicating that CheY Y106 IN states were too short-lived to influence backbone dynamics. CheY*.FliM_N_ clusters had W106 in the IN orientation (θ = 54.1±2.3°). The CheY.FliM_N_ clusters, in striking contrast, spanned the entire Y106 rotamer range. Thus, the intermediate CheY.FliM_N_ Y106 rotamer states were sufficiently stable to affect backbone dynamics (**Figure 6**).

**Figure 6:**
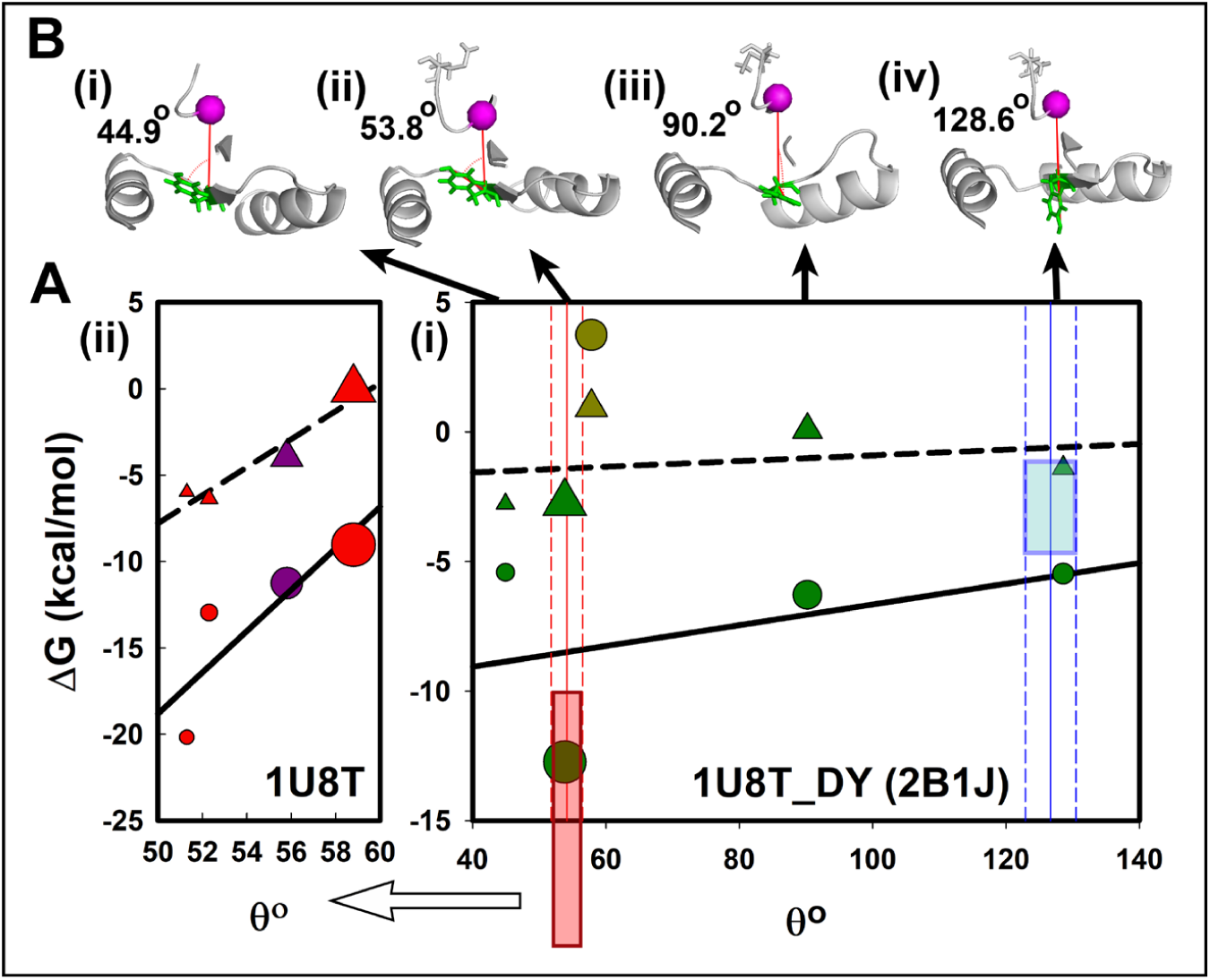
Rotamer Y/W106 energetics. A. Interface and CheY fold stabilization. Interface (ΔG_int_, triangle), CheY fold (ΔG_CheY_, circle). Linear regressions (interfacial (dashed), fold (solid). **(i)** CheY.FliM_N_ (green). 2B1J crystal values (lime). Vertical lines and rectangles show (CheY (cyan) and CheY*. FliM_N_ (red) θ and ΔG_core_ range respectively. Correlations: θ-ΔG_interface_ (R = 0.23, Pearson = 0.63); θ-ΔG_CheY_ (R = 0.43, Pearson = 0.21). **(ii)** CheY*.FliM_N_ (red). 1U8T crystal values (purple). Correlations: θ-ΔG_interface_ (R = 0.96, Pearson = 0.98); θ-ΔG_CheY_ (R = 0.85, Pearson = 0.33) **B.** CheY conformation and Y106 (green) sidechain rotamer orientation in represntatives of the major CheY.FliM_N_ clusters.

Next, we computed the activation energetics by measurement of ionizable residue electrostatics with PROPKA. There was a weak stabilization of the CheY FliM_N_ interface and core with the internalization of the Y106 sidechain, The buried CheY*.FliM_N_ W106 sidechain had a substantially more restricted rotation range than the CheY.FliM_N_ Y106 sidechain. However, the correlation between side-chain orientation and stabilization of CheY* FliM_N_ interface and CheY* core was stronger, consistent with a more-tightly packed CheY* FliM_N_ complex. The stabilization of the interface by the D13K/Y106W residue substitutions was consistent with the different FliM_N_ binding affinities measured in solution for active versus inactive CheY states. The novel result, not recognized previously, was the coupled stabilization of the CheY fold for both CheY.FliM_N_ and CheY*.FliM_N_.

The energetics computed for the 1U8T crystal structure were in line with results from the MD conformational ensembles. In contrast, the values computed for the 2B1J crystal structure were outliers reporting higher energy states relative to the values obtained from the MD runs, an outcome that may be linked to errors in atomic coordinate positions due to the increased B-factor values around the 2B1J CheY.FliM_N_ interface (Supporting Information Figure S2) and/or deformation of the local volume around K109, FIII, M17 by crystal packing contacts (Supporting Information Figure S6).

### 6. An emergent sector orchestrates CheY* allosteric communication

Second, we developed the network model for a comprehensive description of the temporal conformational couplings. The centrality analysis identified network nodes but did not characterize couplings between them with other fragments that constituted (>95%) of the information available in the nMI matrix. We used community analysis (63, 64), to extract and map this information onto the structure. Community networks are collapsed networks that partition and map the protein into semi-rigid bodies (“sectors”) that may be schematized for a concise, comprehensive representation. The schematics and their mapping onto the 3D structure will be henceforth referred to as community “network” and “map” respectively.

Community analysis of native CheY revealed distinct sectors (n > 5) displaying coordinated dynamics. The β3 strand F_53_VISD_57_ occupied a central location in contact with all sectors. Sector A, organized around the D57 phosphorylation site coupled to the other sectors, in particular sector B, organized around the FliM_N_-binding surface. The tCONCOORD CheY* community map, when compared against the corresponding CheY map, showed a small increase in sector A relative to sector C interactions with sector B (**Figure 7A)**. This result may indicate limited activation of CheY* relative to CheY detectible with the more sensitive community versus global network, but does not challenge the conclusion that CheY and CheY* have similar dynamic architecture based on retention the loops as network nodes reinforced

**Figure 7:**
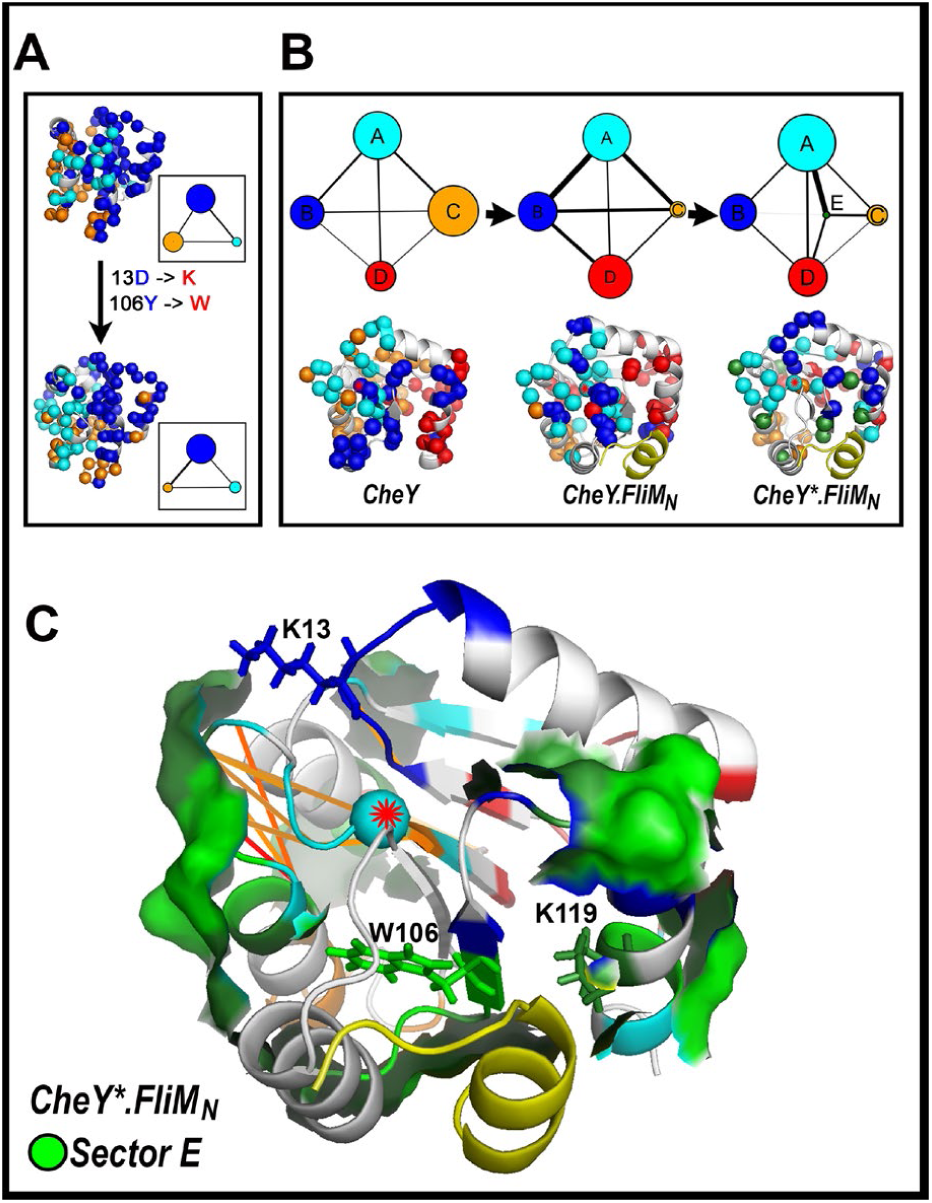
Changes in community network architecture triggered by D13K/Y106W substitutions and FliM_N_ peptide. The reduced number of sectors compared to single fragments as nodes provided a concise, quantitative readout of the protein dynamics. **A. CheY and CheY* community maps.** Networks **(boxed insets)** from tCONCOORD runs show the reduction in the size of sector C relative to sectors A and B in CheY* versus CheY. **B. CheY, CheY.FliM**_**N**_ **and CheY*.FliM**_**N**_ **community architecture.** Networks **(top)** and maps **(bottom).** FliM_N_ = yellow (cartoon representation). The MD detected four dynamic sectors for CheY (A= cyan, B = blue, C = orange, D = red). The sector C from the tCONCOORD runs is resolved into two sectors (C and D) in the MD runs. Node size = sector membership; edge thickness = weighted inter-node interactions). Sectors A and B are built around the phosphorylation site (D57 (red asterisks)) and the FliM_N_ binding surface respectively. They increase at the expense of sectors C upon complex formation. The presence of phospho-mimic mutations in the complex (CheY*.FliM_N_) creates an additional sector E from sectors A and B, that orchestrates interactions with sectors C and D.. **C.** CheY*.FliM_N_ community map showing Sector E surface. See **Supporting Video S5** for 3D perspective. Sidechains identify the mutated residues (K13, W106) and FliM_N_ binding residue K119, a part of Sector E. Sectors are colored as in A. The strength of the top (>+2σ) nMI couplings (lines) couplings are reflected in their thickness and color (low (yellow) -> high (red)). D57 (red asterisk).

The MD resolved the tCONCOORD sector C into two sectors (**Figure 7B**). Importantly, reported residue substitutions partitioned to sectors A and B in the more detailed map according to phenotype (**Table S1**). Positions, where these are known to affect dephosphorylation kinetics (65), mapped to sector A. Residues known to harbor affect rotation bias, suppressor mutations for CW- or CCW-biasing FliM lesions for example (65) mapped to sector B. Positions yielding mutations that affect interaction with the CheY-phosphatase CheZ (66) were adjacent to Sector D, the smallest sector obtained for CheY. Sector C, comparable in size to A, might be expected to influence the overall stability and rigidity of the protein.

Changes in loop dynamics upon complex formation were reflected in the networks (Figure 7B). The couplings between sectors A (phosphorylation) and B (FliM_N_ binding) were strengthened relative to the free protein. Sector B expanded at the expense of sector C and coupled more strongly to sector A in the CheY.FliM_N_ network. The mutated residue K13 was part of a loop that flipped from sector A to sector B. A fifth sector (E (K_45-48_N_62_-L_65-68_A_101_-S_104-107_F_111-114_K_119-123_)) spanned by the mutated residues (K13, W106) formed in the network of the activated-mutant CheY-FliM_N_ complex (CheY*.FliM_N_). The E-sector fragments were drawn from sector A (K_45,_ N_62,_ K_119_), sector B (A_101,_ S_104,_ F_111_) or fragments adjacent to these sectors in the free CheY community network. Sector E formed a surface-exposed ridge that connected the FliM_N_ α-helix, via S_104-107_ and K_119-123_, to sector C residues E_35_ and (via K_45_) E_37_, (**Figure 7C, Supporting Information Video S5)**. The top nMI couplings connected sector E fragments within the central β3-α3 loop to the D57 phosphorylation site. These couplings were unchanged by complex formation.

## Discussion

The results of this study advance our understanding of CheY conformational plasticity and activation in important ways (**Figure 8)**.

**Figure 8:**
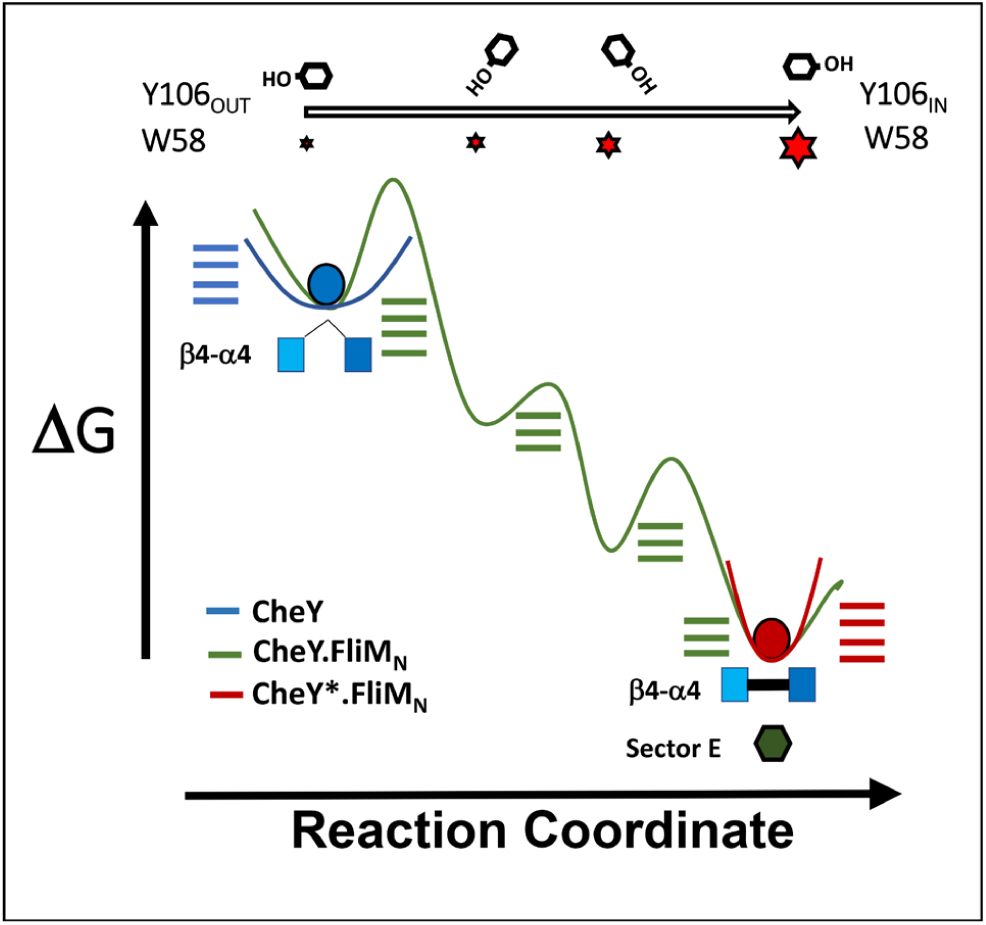
Allosteric priming in E. coli CheY. **A.** Reaction coordinate (x-axis) showing stabilization of the CheY fold coupled with rotamer transitions of CheY aromatic residues Y106 and W58 that have been the common indicators of CheY activation. Horizontal bars indicate multiple local minima. CheY ensembles (blue) have large conformational heterogeneity, controlled by a flexible β4-α4 loop, and sample both Y106 IN and OUT rotamer states; but the IN state is too short-lived to generate CheY sub-populations with distinct backbone conformations. FliM_N_ bound CheY ensembles (green) sample a conformational landscape with a large ΔG range, with prominent troughs among the local minima that track the progressive stabilization of the CheY fold and concerted Internalization of Y106 entrained to tighter FliM_N_ attachment. FliM_N_ bound to D13K-Y106W CheY (CheY*) confines the CheY fold to conformational space (red) around the global minimum. The β4-α4 loop is immobilized by the CheY.K91-FliM_N_ salt-bridge and W106 plus W58 are locked IN a tightly-packed CheY core, with the emergence of a dedicated sector (E) for communication between the phosphorylation site and binding interface. The sector is central to the dynamics of the stabilized CheY core.

### 1. FliM_N_ as an allosteric effector

X-ray crystallography in concert with behavioural and biochemical studies has built a valuable mechanistic framework based on visual inspection of structural landmarks, guided by chemical intuition. Examination of the native CheY.FliM_N_ crystal structure led to the proposal that the complex was an intermediate between active and inactive state consistent with a flexible β4-α4 loop (23). The structure challenged existing two-state switch models; but puzzlingly the central element in the models, the Y106 rotamer state, was not in an intermediate conformation but the activated rotamer state and the decrease in FliM_N_ affinity relative to the activated complex was difficult to discern. These issues have been resolved by the MD simulations and XFMS measurements reported in this study.

The CheY.FliM_N_ conformational landscape generated by MD simulations of the reverse-engineered 1U8T_DY structure had prominent minima that reflected intermediate FliM_N_ attachment entrained to Y106 rotation states that ranged between the dominant OUT state in free CheY and the W106 IN state in activated CheY*.FliM_N_. XFMS determined solvent accessibility values for the CheY.FliM_N_ allosteric relay sidechains that were intermediate between values obtained for inactive CheY and active CheY*.FliM_N_. These values were correlated with the protection of the interfacial lysine residues that monitored FliM_N_ attachment. The D13K-Y106W residue substitutions as seen in the crystal structures did not alter the CheY fold to any significant extent in the absence of FliM_N_; a result supported in this study by both simulation and measurement. The MD clarified that FliM_N_ stabilized CheY and strengthened allosteric communication between its binding interface and the D57 phosphorylation site due to formation, in part, of the CheY.K91-FliM_N_.D3 salt-bridge. The salt-bridge decreased the flexibility of the β4-α4 hinge, consistent with earlier studies (23, 33).

### 2. The dynamics and energetics of activation

This study documents a broad, high-energy CheY conformational landscape with shallow minima consistent with the high conformational plasticity suggested by the CheY crystal structures and early MD studies (Introduction). Network analysis, based on mutual information between short protein fragments established that two loops (β3-α3, β4-α4) act as flexible hinges to control the dynamics. The CheY MD trajectories revealed episodes where the Y106 sidechain is buried (IN), but cluster analysis determined the inward motions were too brief to influence backbone dynamics in contrast to the case for CheY.FliM_N_. The buried states of the Y106 sidechain have not been visualized to our knowledge in inactive CheY crystal structures.

The CheY conformations of the major CheY.FliM_N_ clusters were more stable than the dominant CheY conformations reported by the MD or the conformation in the 2B1J crystal structure. The lifetimes of the CheY Y106 IN states in CheY.FliM_N_ were substantially greater than in free CheY and represented in the major clusters, There was a weak correlation between the stability of the CheY fold, the FliM_N_ interface and the position of the Y106 sidechain. The CheY ΔG values in the major CheY.FliM_N_ clusters overlapped with the values in the inactive CheY and activated CheY*.FliM_N_ clusters.

The mean CheY*.FliM_N_ ΔG value was more stable than for CheY.FliM_N_. This was also the case for the interfacial ΔG values. The position of the W106 sidechain was restricted to a narrow range. Nevertheless, the ΔG values for both the CheY fold and its FliM_N_ interface, as well as the rotamer positions of the W106 and Y106 sidechains, were similar for the most dominant CheY*.FliM_N_ and CheY*.FliM_N_ conformational clusters. The similarity may explain capture during crystallization of the Y106 sidechain in the 2B1J structure in a position superimposable with the W106 sidechain in the 1U8T structure. The better correlation of W106 sidechain position, in the MD clusters and the 1U8T structure, with the CheY fold and FliM_N_ interface ΔG values, reflects the tight-packing due to the D13K-Y106W substitutions. The ΔG and W106 rotation angle values of the CheY*.FliM_N_ clusters had no overlap with values for CheY clusters.

Allosteric communication may range from largely enthalpic, as in lysozyme, to largely entropic, with the change in flexibility rather than shape (67). Both energy terms contribute to CheY allosteric activation. CheY activation has aspects that “invoke conformational selection”, namely the selection of the global minimum from the multiple minima sampled by the native CheY.FliM_N_ conformational ensemble by the D13K-Y106W residue substitutions. Other aspects, such as the formation of the allosteric relay based on local changes in the loop and sidechain rotamer dynamics triggered by FliM_N_ attachment support “induced fit”. Neither description is complete.

### 3. Community networks – a new measure for response regulator signal transduction

It has long been recognized that two-state allosteric models have heuristic value but that a more complete analytical description is desirable (24). The number of states multiplies to accommodate the dynamics (68) as characterization becomes more detailed. Recent work on the multiple motile responses triggered by CheY homologs even within one species (e.g. *Caulobacter crescentus* (69)), as well as the variable signal transduction strategies employed by response regulators (1), have emphasized the need for a more complete description. Community networks have been used previously (64) to identify jointly moving regions that do not track backbone secondary structure but are frequently governed instead by side-chain motions. This work is the first application of this approach to the response regulator superfamily.

Distinct protein sectors with correlated motions were identified in community networks. The extensive library of CheY residue substitutions was exploited for functional assignment of the sectors. Two sectors, namely the neighbourhood of the phosphorylation site (sector A) and the region of FliM_N_ binding (sector B) had clear functional importance. Two other sectors lacked strong, specific phenotypes and might have broader functions in maintaining the overall CheY fold. The long β3-α3 loop influenced movements of the β3 strand that formed a sector junction, consistent with its central role in the reported PC motions. Similar motions take place in other proteins that utilize β-sheets for signal transduction (70).

FliM_N_ attachment increased the size of sectors A and B in the CheY community network. The CheY*.FliM_N_ community network was distinguished from the CheY and CheY.FliM_N_ networks by a fifth sector (E), drawn from sectors A and B, that formed a dedicated conduit between the phosphorylation and FliM_N_-binding sites to cement the allosteric linkage, with the substituted residues K13 and W106 at its boundaries. Sector E in addition to increasing FliM_N_ affinity may influence the regulation of phosphorylation by proteins whose binding surface overlaps with FliM_N_ (68). The emergence of sector E was tied to the closure of the β4-α4 hinge by the CheY.K91-FliM_N_.D3 salt-bridge and “freeze-out” of W_106_VVKP β-strand dynamics by the burial of aromatic residues for a tightly packed core. This sector connects with all other sectors and has a large surface profile. It may directly or indirectly define a region important for binding to FliN.

Rotamer reorientation of aromatic sidechains is a common theme in phospho-proteins, but diverse strategies for coupling side-chain motions to phosphorylation exist. In eukaryotic protein kinases, activation is controlled by DFG motif loops. These loops take on multiple IN and OUT orientations, with orientation correlated with activation. In Aurora kinase A, phosphorylation triggers transition between distinct IN orientations, rather than between IN and OUT states (71). In calcium calmodulin-dependent kinase, IN and OUT DFG states are loosely coupled to kinase domain phosphorylation (72). in CheY XFMS reported D_57_WN_59_ internalization was coupled to protection at the FliM_N_ interface. We envisage that XFMS will have applications in other phosphorelays given ongoing developments in MS sensitivity and high-throughput analyses since most amino acids are modified by hydroxy radicals to a greater or lesser extent

The sparse sampling by crystal structures may miss high-energy states, such as the intermediate states of the CheY 106 sidechain, that are important for deciphering mechanism. MD simulations provide a much more detailed sampling of the conformational landscape, but their challenge is to extract the essential features from the large conformational ensembles obtained; a challenge only partially met by standard PCA and RMSF analyses. Our study shows that community maps provide a concise, comprehensive description based on quantitative criteria for identification of the key features of CheY allosteric activation, to make the case that they provide the optimal compromise for mechanistic dissection of signal transduction strategies in the response regulator superfamily.

## AUTHOR CONTRIBUTIONS

**Paige Wheatley:** Investigation, Visualization, Writing - Review & Editing

**Sayan Gupta:** Methodology, Investigation

**Alessandro Pandini:** Conceptualization, Software, Formal analysis, Validation, Visualization

**Yan Chen:** Methodology, Investigation, Formal analysis, Validation

**Christopher J Petzold:** Methodology, Supervision, Funding Acquisition, Writing - Review & Editing

**Corie Y. Ralston:** Conceptualization, Supervision, Funding Acquisition, Writing - Review & Editing

**David F. Blair:** Conceptualization. Supervision, Funding Acquisition, Writing - Draft, Review & Editing

**Shahid Khan:** Conceptualization, Formal analysis, Writing - Draft, Review & Editing, Project Administration

## ACKNOWLEDGEMENTS

We thank Dr. Robert Bourret for comments on the manuscript and Martin Horvath for assistance with the FPLC analysis of CheY proteins. This study was supported by National Institutes of Health grants 1R01GM126218 (to C.Y.R) and R01GM46683 (to D.F.B.). The XFMS was conducted at the Advanced Light Source beamline 3.2.1 and the Joint BioEnergy Institute, supported by the Office of Science, Office of Biological and Environmental Research, of the U.S. DOE under contract DE-AC02-05CH11231. The MD simulations described in this paper were executed on the Crick Data Analysis and Management Platform (CAMP), provided by the Francis Crick Institute. Other computations utilized the Molecular Biology Consortium computer cluster.

