## Supporting Information for "Allosteric priming of *E. coli* CheY by the flagellar motor protein FliM"

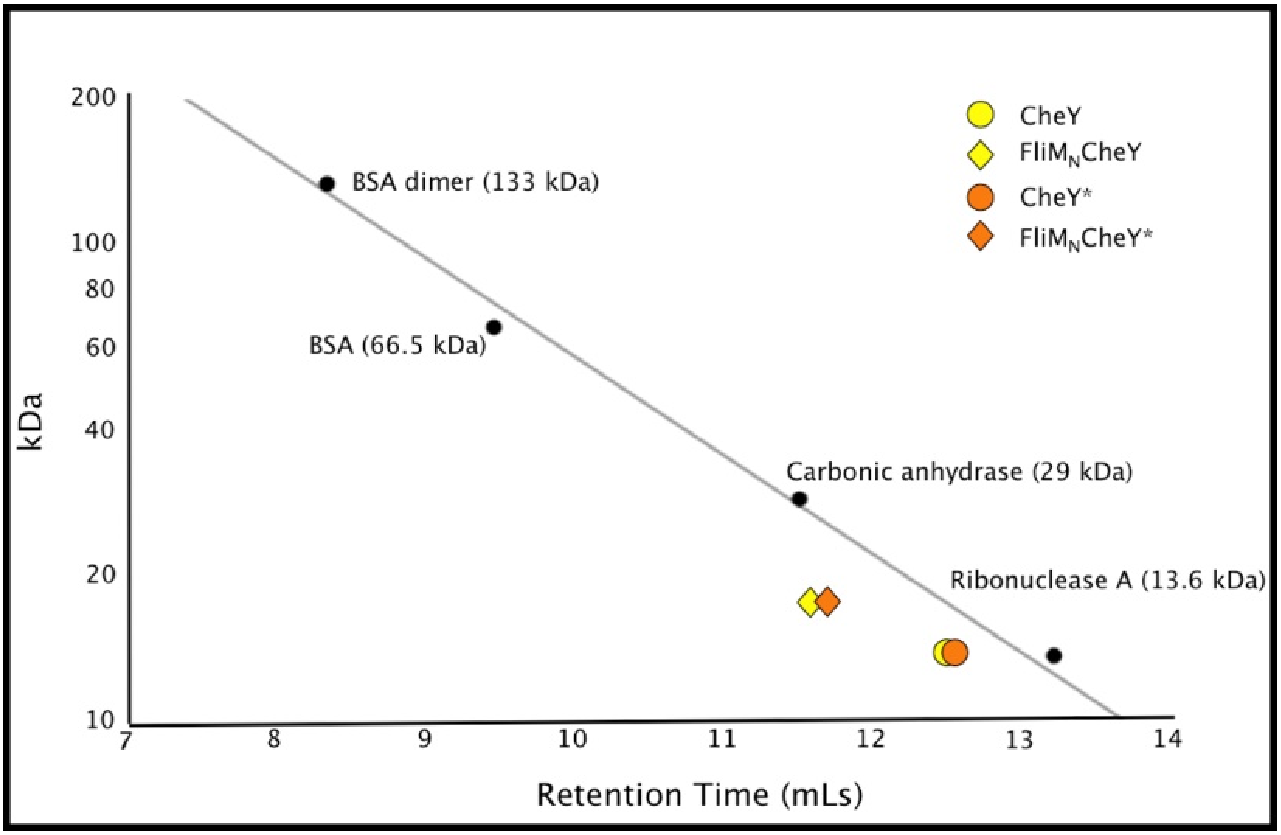


**Figure S1: FPLC analysis of purified CheY proteins.** *FPLC of purified CheY, CheY D13K Y106W (CheY*), FliM$_N$CheY, and FliM$_N$CheY* was performed on an AKTA Superdex 75 10/300 GL column in XF buffer to determine whether the purified proteins were in a monomeric state. BSA, carbonic anhydrase, and ribonuclease A were used as molecular weight standards. The calculated molecular weights of CheY, CheY*, FliM$_N$CheY, and FliM$_N$CheY* are 15.3 kDa, 15.3 kDa, 18.5 kDa, and 18.5 kDa, respectively. The apparent molecular weights of CheY, CheY*, FliM$_N$CheY, and FliM$_N$CheY* are 17.5 kDa, 16 kDa, 28.5 kDa, and 27.5 kDa, suggesting that each purified protein is monomeric in solution.* $R^2 = 0.99$

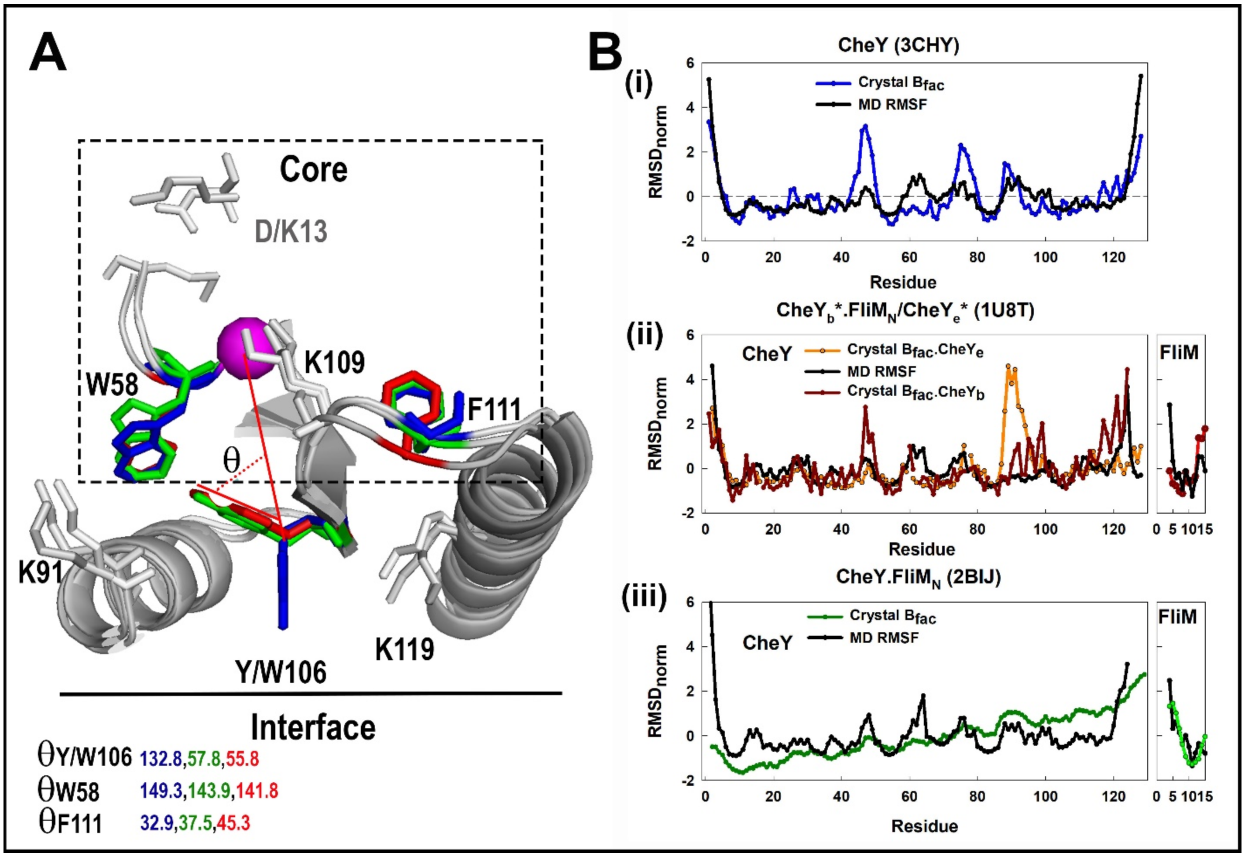


**Figure S2:** ***A. Crystal structures rotamer orientations.*** *The structures are colour-coded (3CHY (blue), 2B1J (green), 1U8T (red)). Sidechain orientation ($\theta$) of the interfacial residue Y/W106, as well as the core residues W58 and F111. The orientation angle $\theta$ was measured between the line connecting the residue (R) and D57 $C^{\alpha}$ (magenta sphere) atoms and the line connecting R $C^{\alpha}$ with the most distant heavy atom in the aromatic sidechain (OH (Y), CZ (F), CZ2 (W))). Inward rotation of the residues is correlated. The $\theta$s in the 2B1J structure have intermediate values between values for 3CHY and 1U8T. The deviation in $\omega$ did not alter the correlation. For 1U8T D13K-Y106WCheY: $\theta$ Y106 = 130.0$^{o}$, $\theta$ W58 = 160.4$^{o}$, $\theta$ F111 = 32.9$^{o}$.* ***B. $C^{\alpha}$ backbone dynamics - MD versus crystal structures.*** *Root-mean square fluctuations (RMSFs) from MD simulations compared against X-ray crystal B-factors ($B_{fac}$). The normalized profiles $((X_{i=0->n} - (\Sigma X/n)/\sigma_X)$ are shown. Mean RMSF for CheY.FliM$_N$ (2BIJ) was not stationary due to upward drift in the C-terminal part of CheY that forms the FliM binding surface and FliM$_N$.*

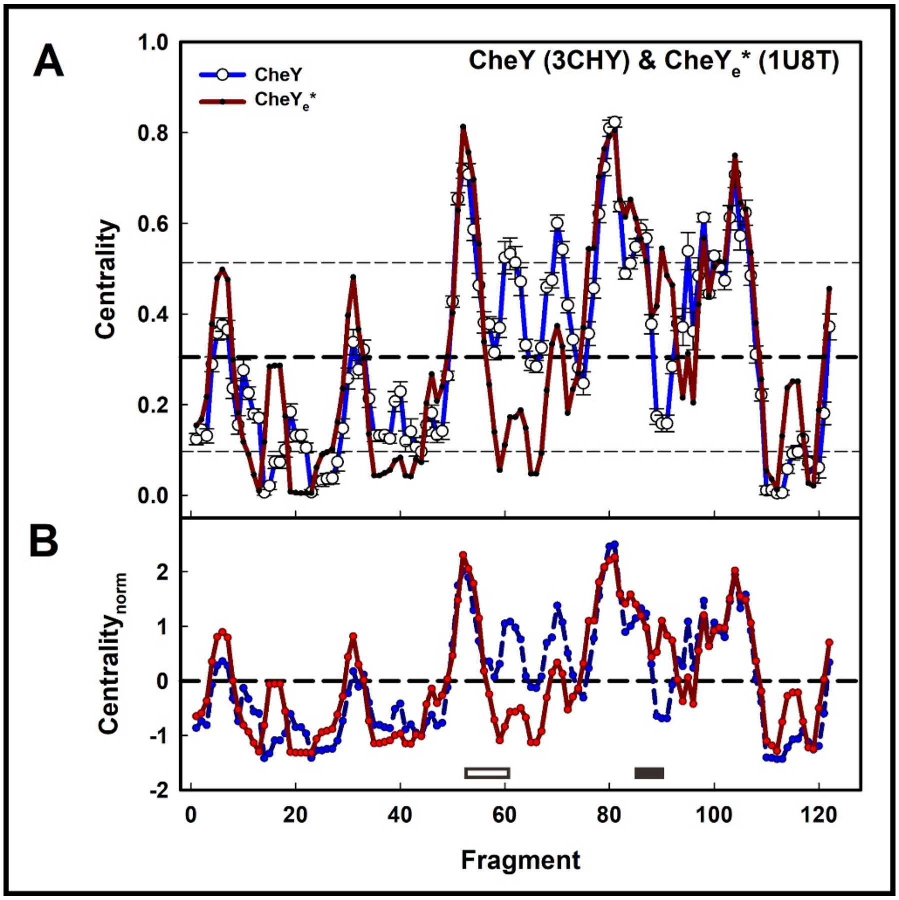


**Figure S3: Mutation induced changes in C$^{\alpha}$ backbone dynamics.** *Comparison of the dynamics of CheY and mutant (13DK106YW) CheY (CheY*) with tCONCOORD [1]. Crystal structures from which the ensembles were derived are bracketed.* **A.** *Eigencentrality profiles. Mean (dashed line)* $\pm$ $\sigma$ *(dotted lines). Fragment number = 1st residue of a 4-residue fragment.* **B.** *Normalized* $((X_{i=0\text{->}n} - (\Sigma X/n)/\sigma_X)$ *eigencentrality profiles. Zero (dashed line). CheY* $\theta$*Y106 = 131* $\pm$ *3°. CheY** $\theta$ *W106 = 130* $\pm$ *3°. Horizontal bars indicate loops as in Figure 2B.*

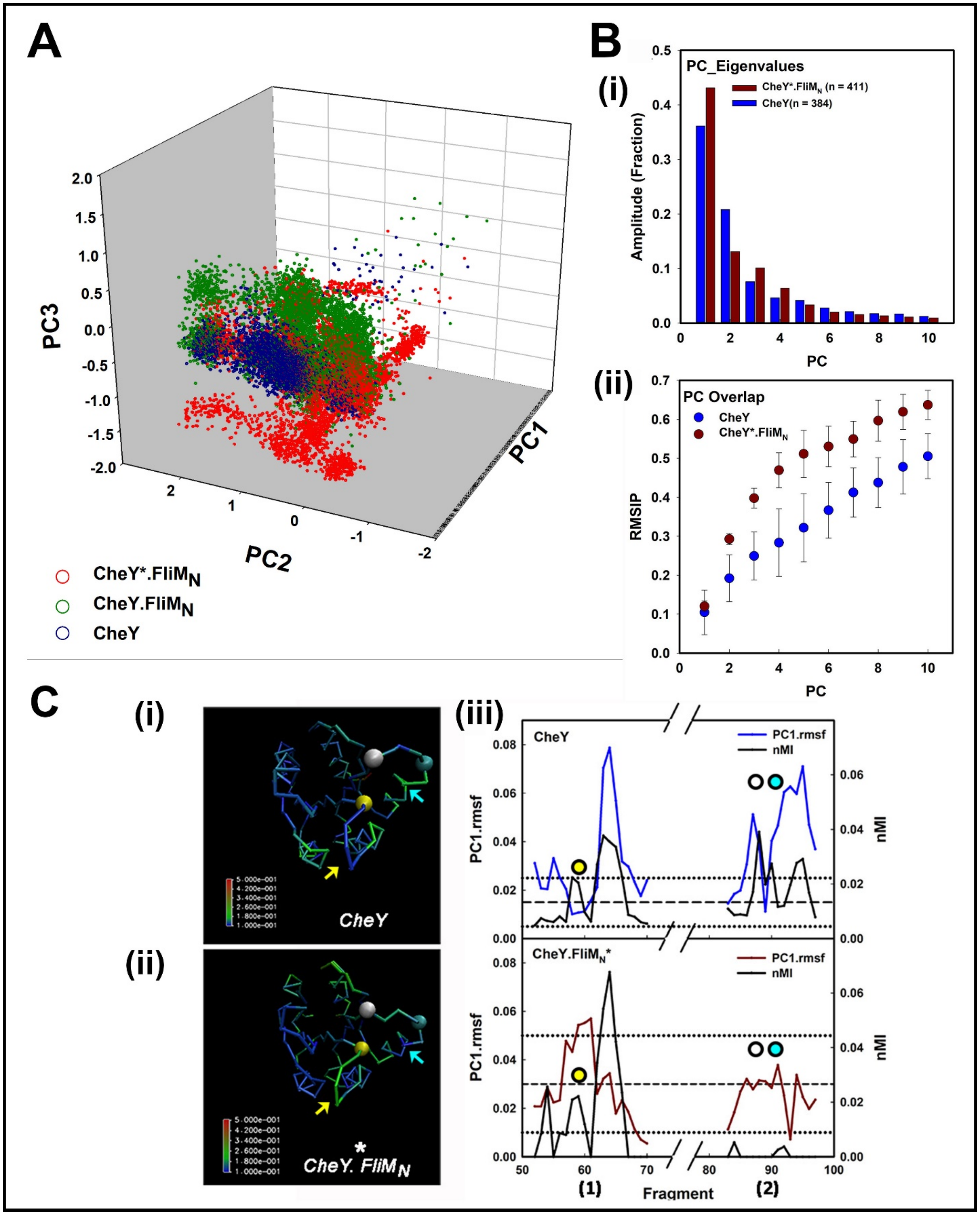
A
B
(i)
PC_Eigenvalues
CheY*.FliMN (n = 411)
CheY(n = 384)
Amplitude (Fraction)
PC
(ii)
PC Overlap
CheY
CheY*.FliMN
RMSIP
PC
PC3
PC1
PC2
CheY*.FliMN
CheY.FliMN
CheY
C
(i)
(ii)
(iii)
CheY
CheY. FliMN
CheY.FliMN*
PC1.rmsf
nMI
Fragment
(1)
(2)

**Figure S4: Conformational ensembles populated by FliM peptide and activating mutations**. ***A. PC1-PC2-PC3 plot.*** *The MD trajectories of the three structures (CheY, CheY.FliM$_N$, CheY*.FliM$_N$) were merged based on a common CheY atom set. The member structures were projected onto the first three PCs. Non-overlapping structures indicate new conformational space made accessible by the binding of FliM$_N$. Distinct CheY conformational ensembles are sampled by CheY, CheY.FliM$_N$, and CheY*.FliM$_N$.* ***B. (i) Eigenvalue amplitudes for the first 10 PCs.*** *The first 3 PCs account for about 60% (CheY) and 70% (CheY*.FliM$_N$) of the total amplitude of all PCs (=n). The PC1 amplitude for both CheY and CheY*.FliM$_N$ accounts for about 40% of the total amplitude.* ***(ii).*** *Evaluation of the root mean square inner product (RMSIP) establishes that the essential conformational sub-space is captured by the first 10 PCs over all replica runs for each structure.* ***C. Hinge characterization.*** *PC1 RMSFs for their complete MD ensembles were mapped onto the structures.* ***(i).*** *CheY (3CHY)*. ***(ii).*** *CheY*.FliM$_N$ (1U8T). Colour-coded bars indicate RMSF values. Spheres denote W58 (yellow), K91 (cyan) and T87 (white) residues. Arrows identify $\beta$3-$\alpha$3 (yellow) and $\beta$4-$\alpha$4 (cyan).* ***(iii).*** *PC1 superimposed fragment nMI and RMSF values for the loops $\beta$3-$\alpha$3 (1) and $\beta$4-$\alpha$4 (2). The horizontal reference lines are CheY PC1 RMSF (mean (dashed)$\pm\sigma$ (dotted)) values. The values are 0.015$\pm$0.01(3CHY); 0.03$\pm$0.02 (1U8T). Fragments with W58, T87 and T91 are marked (circles colour-coded as in A, B).*

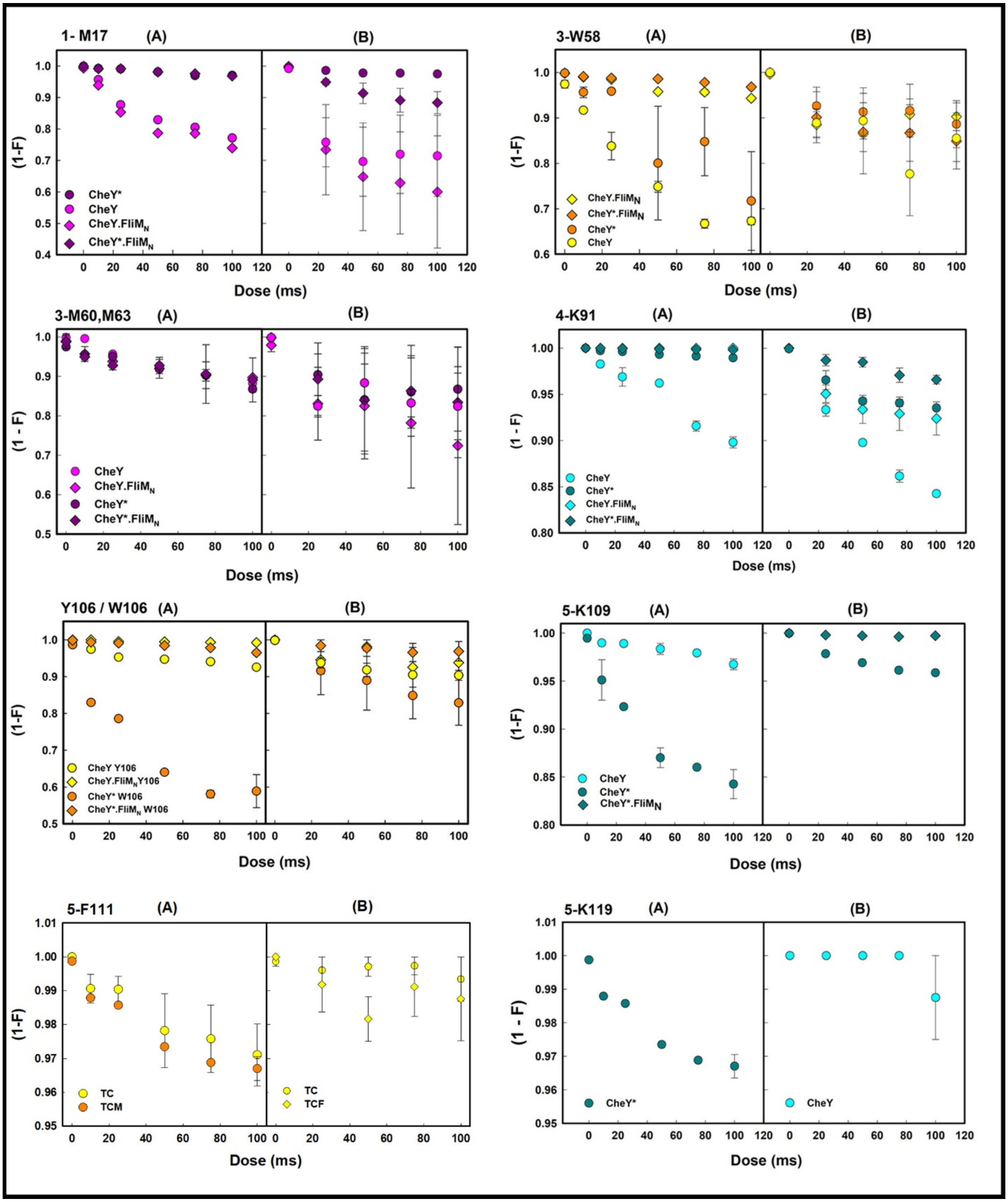
1- M17 (A) (B)
CheY*
CheY
CheY.FliMN
CheY*.FliMN
3-W58 (A) (B)
CheY.FliMN
CheY*.FliMN
CheY*
CheY
3-M60,M63 (A) (B)
CheY
CheY.FliMN
CheY*
CheY*.FliMN
4-K91 (A) (B)
CheY
CheY*
CheY.FliMN
CheY*.FliMN
Y106 / W106 (A) (B)
CheY Y106
CheY.FliMNY106
CheY* W106
CheY*.FliMN W106
5-K109 (A) (B)
CheY
CheY*
CheY*.FliMN
5-F111 (A) (B)
TC
TCM
TC
TCF
5-K119 (A) (B)
CheY*
CheY
(1-F)
(1 - F)
Dose (ms)

**Figure S5: Complete XFMS data for the examined residues**. *Two technical replicates from 2 independent experiments (A, B) were analyzed. Residues were selected from a larger set based on high intrinsic reactivity (Table 1 of [2]). The reactivity followed the order – M, F, Y, W, R, K (high -> low). Oxidation of W106 in CheY* was greater than that of CheY Y106 consistent with X-ray crystallographic evidence [3] that W106 is exclusively OUT in CheY*. The increase may be due, in part, to greater intrinsic reactivity (W/Y=1.45) and/or difficulty in packing the bulkier tryptophan within the protein core in the absence of FliM$_N$ binding free energy. The lysines (K91, K109, K119) participate in ionic or hydrogen bonds. K91 and K119 form salt-bridges with FliM$_N$ D3 and D12 respectively in CheY*.FliM$_N$. These bonds are weaker in CheY.FliM$_N$, consistent with the crystal structures [3]. K109 hydrogen-bonding reported by the crystal structures (Introduction) was noted, but not analyzed in our MD simulations. There is no detectible K119 oxidation in the fusion proteins, but the lower rate of K91 oxidation in CheY*.FliM$_N$ versus CheY.FliM$_N$ supports the MD. K109 oxidations in the fusions were very limited (FliM$_N$.CheY*) or undetectable (FliM$_N$.CheY). The lack of K119 and K109 oxidation in the fusions is difficult to explain solely by sidechain burial, since aromatic residues W106 or W58 have detectible, albeit reduced, oxidation in FliM$_N$.CheY*. Bonding interactions may contribute by conferring resistance to hydroxy radical attack.*

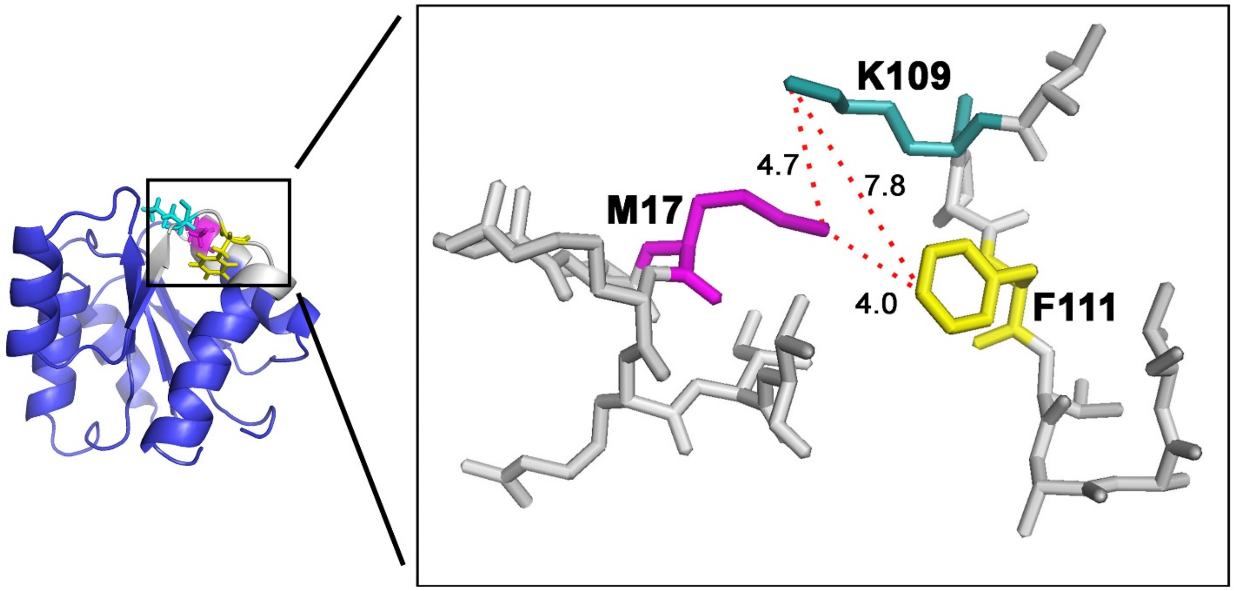


**Figure S6**: **Anomalous oxidation is localized to a small CheY cluster.** *Nearest atom distances (in angstroms) between the residues M17, K109 and F111 (stick representations) are shown (dotted red lines). Outliers beyond the 95% confidence limit (Figure 9) were found exclusively for cases involving these residues. Weighting with the intrinsic reactivity [2] did not significantly improve alignment with the overall SASA-log(PF) relation. SASA weighting by a contact parameter [4] also did not improve its correlation with log(PF) [2]. The crystal structures may not represent the solution conformation of this loop. Alternatively, hydroxy radical reactions could be influenced in ways that are not presently fully understood. Methionines are susceptible to secondary solvent reactions [5]. Local perturbation of the bound water in the adjacent phosphorylation site, caused by a disrupted hydrogen-bonding network reported for the CheY D13K crystal structure [6], may affect oxygen radical reactivity of these sidechains [7]; in addition to bonding interactions, their alteration [6] or modulation by the local dielectric. Future MD simulations are being designed to analyze bonding interactions and their effect of oxygen radical reactivity.*

| RESIDUE | SECTOR | MUTATION | CITATION |
|---|---|---|---|
| D12 | A | D -> E, N | [8] |
| D13 | A | D -> E, N, K | [6, 8, 9] |
| S56 | A | S -> F | [10] |
| N59 | A | F -> * | [11] |
| E89 | A | F -> * | [11] |
| D57 | A/B | D -> E, N | [8, 9] |
| T87 | B | T -> I | [9, 12, 13] |
| A90 | B | FliM$^{sup}$, A->V | [10] |
| K91 | B | K -> R. Ac | [14] |
| K92 | B | K -> R. Ac | [15] |
| I95 | B | I -> V | [16] |
| Y106 | B | Y -> W | [12] |
| K109 | B | K -> R. Ac | [9, 14] |
| V108 | B$^{\#}$ | FliM$^{sup}$, V->M | [10] |
| F111 | B$^{\#}$ | FliM$^{sup}$, F->V | [10] |
| T112 | B | FliM$^{sup}$, T->I | [10] |
| F14 | C | F -> * | [11] |
| E27 | D | FliM$^{sup}$, E->K | [10] |
| K26 | D$^{\#}$ | K -> E | [17] |
| E117 | # | FliM$^{sup}$, E->K | [10] |
| N23 | # | N -> D | [17] |

**Table S1: Mapping of residue substitutions to sectors in the CheY community map.** *Asterisks denote cases where the substituted residue does not map to a sector. Superscripted asterisks mark sectors < 3 positions (< 1 fragment) removed from an unassigned mutated residue. Ac = Acetylation affects chemotaxis.*

**Movie S1: CheY dynamics.** *The raw trajectory for 3CHY showing movements of the Y106 side chain (red) and T87 (green)*

**Movie S2: Dynamics of the inactive CheY.FliM$_N$ complex.** *The raw trajectory for the native complex (1U8T_DY), engineered from 1U8T, showing movements of the Y106 (red) and T87 (green) side chains).*

**Movie S3: Dynamics of the activated mutant CheY*.FliM$_N$ complex.** *The raw trajectory for 1U8T, showing movements of the W106 side chain (red) with T87 (green).*

**Movie S4: CheY.FliM$_N$ interface dynamics.** *Interface extracted from the raw trajectory of the 1U8T_DY complex. Sidechains (CheY 106Y (red) and K119 green), FliM$_N$ D12 (pink)).*

**Movie S5: Sector coupling the FliM$_N$ interface with the phosphorylation site.** *The CheY*.FliM$_N$ community map showing the surface profile of the coupling sector E (dark-green, green (adjacent residues)) specific for CheY*.FliM$_N$. Top nMI couplings (lines (orange (low) -> red (high)) connect sector E to D57. The $\beta$3 strand forms a junction for sectors A (red), B (orange) and C (cyan). Sector D (blue). Sidechains (K13, D57, T87, W107) are colored according to their sector affiliation.*

-----------------------------------------------------------------------------------------------------
